# Long G4-rich enhancer physically interacts with EXOC3 promoter via a G4:G4 DNA-based mechanism

**DOI:** 10.1101/2024.01.29.577212

**Authors:** Jeffrey D DeMeis, Justin T Roberts, Haley A Delcher, Noel L Godang, Alexander B Coley, Cana L Brown, Michael H Shaw, Sayema Naaz, Enas S Alsatari, Ayush Dahal, Shahem Y Alqudah, Kevin N Nguyen, Anita D Nguyen, Sunita S Paudel, Hong Dang, Wanda K. O’Neal, Michael R. Knowles, Dominika Houserova, Mark N Gillespie, Glen M Borchert

## Abstract

Enhancers are genomic sequences that function as regulatory elements capable of increasing the transcription of a given gene often located at a considerable distance. The broadly accepted model of enhancer activation involves bringing an enhancer-bound activator protein complex into close spatial proximity to its target promoter through chromatin looping. Equally relevant to the work described herein, roles for guanine (G) rich sequences in transcriptional regulation are now widely accepted. Non-coding G-rich sequences are commonly found in gene promoters and enhancers, and various studies have described specific instances where G-rich sequences regulate gene expression via their capacity to form G-quadruplex (G4) structures under physiological conditions. In light of this, our group previously performed a search for long human genomic stretches significantly enriched for minimal G4 motifs (referred to as LG4s herein) leading to the identification of 301 LG4 loci with a density of at least 80 GGG repeats / 1,000 basepairs (bp) and averaging 1,843 bp in length. Further, in agreement with previous reports indicating that minimal G4s are highly enriched in promoters and enhancers, we found 217/301 LG4 sequences overlap a GeneHancer annotated enhancer, and the gene promoters regulated by these LG4 enhancers were found to be similarly, markedly enriched with G4-capable sequences. Importantly, while the generally accepted model for enhancer:promoter specificity maintains that interactions are dictated by enhancer- and promoter-bound transcriptional activator proteins, the current study was designed to test an alternative hypothesis: that LG4 enhancers physically interact with their cognate promoters via a direct G4:G4 DNA-based mechanism. As such, this work employs a combination of informatic mining and locus-specific immunoprecipitation strategies to establish the spatial proximity of enhancer:promoter pairs within the nucleus then biochemically confirms the ability of individual LG4 ssDNAs to directly and specifically interact with DNA sequences found in their target promoters. In addition, we also identify four single nucleotide polymorphisms (SNPs), occurring within a LG4 enhancer on human chromosome 5, significantly associated with Cystic Fibrosis (CF) lung disease severity (avg. p value = 2.83E-9), presumably due to their effects on the expressions of CF-relevant genes directly regulated by this LG4 enhancer (e.g., EXOC3 and CEP72).

**Graphical Abstract:** 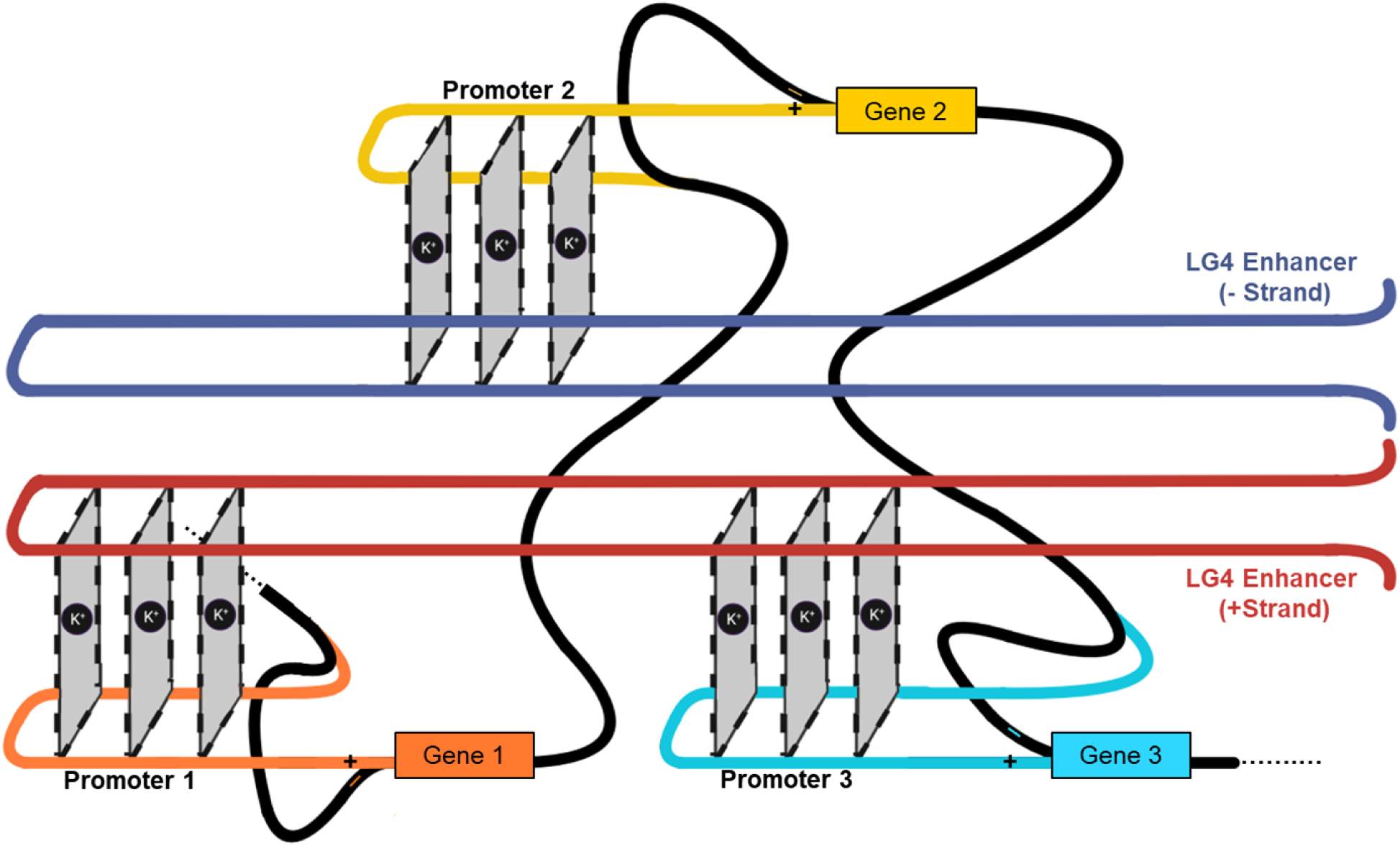

In brief: LG4 enhancers physically interact with gene promoters by forming composite G4 structures where both the LG4 and cognate promoter contribute half of the necessary sequence for G4 formation.

## Introduction

Enhancers are genomic sequences that function as cis-acting regulatory elements capable of increasing the transcriptional abundance of a given gene(s) by recruiting transcription factors and RNA polymerase to a target promoter^1^. Enhancers are often located at a considerable distance from target promoters, and the broadly accepted model of enhancer activation is that an activator protein complex assembles on an enhancer before being transferred to the target promoter^2^. First proposed in prokaryotes in 1986, the looping model of enhancer:promoter communication^3^ (in which looping of the DNA separating an enhancer and its target promoter brings these regulatory elements into close proximity) is now almost universally accepted^1^. Notably, the development of methods aimed at elucidating the spatial organization of the genome via proximity ligation has helped firmly establish this mechanism as the predominant form of enhancer:promoter communication in vertebrates^4^. As an example, a recent micro-C analysis confirmed that more than 65% of experimentally verified enhancer:promoter pairs associate through enhancer:promoter looping in human K562 lymphoblast cells^5^. In addition, the ability of enhancers and promoters to contact one another is also partially controlled through organizing chromosomes into topologically associated domains which often directly limits the range of enhancer influence^6–9^. That said, enhancers frequently demonstrate selectivity for specific promoters within individual topologically associated domains with enhancers routinely crossing other intervening genes that are not activated by the enhancer during the “search” for a target promoter^2,10,11^. Several models have been proposed to explain promoter selectivity, but to date, the mechanisms responsible for the majority of specific enhancer:promoter interactions remain unresolved.

Equally relevant to the work described herein, roles for guanine rich repetitive sequences in transcriptional regulation are now widely accepted^12–14^. Non-coding guanine (G) rich sequences are commonly found in gene promoters and enhancers, and several independent studies have described specific instances where G-rich repetitive sequences regulate gene expression via their capacity to form transient G-quadruplexes (G4s) under physiological conditions^15^. G4s are non-B form DNA secondary structures that arise from Hoogsteen base pairing between G nucleotides stabilized by a central cation^16^ (**Figure 1A**). While there is a large degree of heterogeneity among sequences capable of forming G4s, the minimally defined sequence thought to be required for G4 formation consists of four adjacent runs of G triplets separated by a spacer sequence of varying length (GGGnGGGnGGGnGGG)^16,17^ (**Figure 1B**).

**Figure 1.**
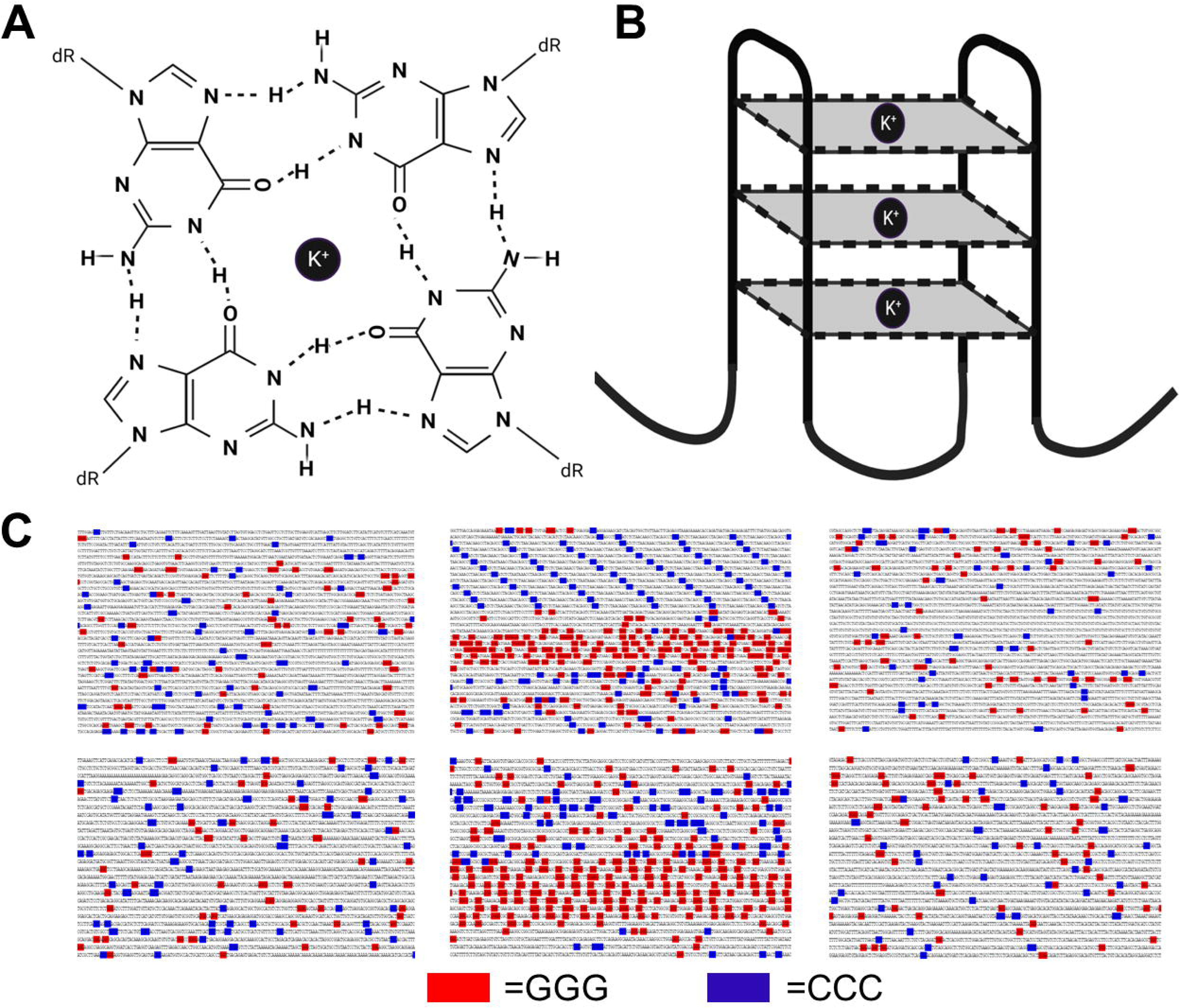
G4 DNA. (**A**) Structure of an individual G-quartet wherein guanine nucleotides are held together by hydrogen bonds centered about a central potassium cation (K+). (**B**) Cartoon depiction of unimolecular antiparallel G4 DNA where each square corresponds to an individual G-quartet and the corners of each quartet correspond to a single guanine. (**C**) Two examples of LG4 loci with ≥3 consecutive genomic Gs shown in red and ≥3 consecutive genomic Cs shown in blue . Top center, LG4 located at human GRCh38 Chr5:551935:556936:1; Top left and right, ∼5kb sequences located 100kb upstream and downstream of the central LG4 (GRCh38:5:451935:456936:1 and GRCh38:5:651936:656935:1 respectively); Bottom center, LG4 located at human GRCh38 Chr12:132686134:132690031:1; Bottom left and right, ∼5kb sequences located 50kb upstream and downstream of the central LG4 (GRCh38:12:132636134:132640031:1 and GRCh38:12:132736134:132740031:1 respectively).

Search algorithms modeled after variations of this minimal G4 motif routinely identify hundreds of thousands of putative G4 capable sequences in the human genome^18–20^, although the true number of G4 forming loci, as well as the timing and cellular circumstances during which G4 structures actually form, have yet to be fully described.

In light of this, instead of focusing on shorter, more rigidly defined G4 motifs, our group previously performed a study in which we searched the human genome for long genomic stretches significantly enriched for minimal G4 motifs (referred to as LG4s herein). Since G-rich immunoglobulin switch regions^21,22^ are composed of a high percentage of (and routinely form) G4 structures and are characteristically associated with DNA breaks and altered gene expressions^21,22^, we elected to model our search parameters on the G-rich Sµ immunoglobulin switch region successfully leading to the identification of 301 LG4 loci with a density of at least 80 GGG repeats / 1,000 basepairs (bp) and averaging 1,843 bp in length (see examples in **Figure 1C**)^23^. Further, in agreement with previous reports indicating that minimal G4s are highly enriched in promoters and enhancers^13,14^, we found LG4s (in comparison to size matched control loci) significantly associated with transcriptional regulatory elements^24,25^ and enriched in 26 different transcription factor (TF) ChIP-seq datasets^26,27^ (including CTCF which is known to facilitate enhancer:promoter interactions and long-distance enhancer-dependent transcription^28^). Furthermore, and of particular relevance to this study, we also found 217/301 LG4 sequences overlap a GeneHancer^29^ annotated enhancer, and our initial analyses of the gene promoters regulated by these LG4 enhancers (as indicated by GeneHancer) found these promoters similarly, markedly enriched with G4-capable sequences^23^.

Importantly, while the current generally accepted model for enhancer:promoter communication involves initial targeting of specific sequences in genomic enhancers and promoters by transcriptional activator proteins followed by enhancer:promoter engagement being facilitated by interactions between these proteins^1,11^, this study was designed to test an alternative hypothesis: that LG4 enhancers physically interact with their cognate promoters via a direct G4:G4 DNA based mechanism. As such, in this work we employ a combination of informatic mining and locus-specific immunoprecipitation strategies to establish the spatial proximity of interacting loci within the nucleus then biochemically assess the ability of specific genomic DNA sequences to interact with one another directly.

## Results

### Chr 5 and Chr 12 LG4s are enhancers annotated as regulating G4 enriched promoters

After finding 217 of the 301 LG4 loci identified in our initial study either fully or partially overlap with an annotated human enhancer (versus only 84 average overlaps (n = 5) between size and nucleotide composition matched control loci and enhancers)^23^, we decided to select two LG4 loci (independently annotated as high confidence enhancers by multiple groups) for experimental validation. Notably, the first of theseLG4s located at human chromosome 5p15.33 (GRCh38:5:551935:556936:1) directly overlaps 13 reported enhancers independently annotated in the Ensembl regulatory build^24^, the Encyclopedia of DNA Elements (ENCODE)^30^, GeneHancer^29^, and SuperEnhancer^31^ datasets. Similarly, we also selected a second LG4 located at human chromosome 12q24.33 (GRCh38 Chr12:132686134:132690031:1) which directly overlaps 9 reported enhancers independently annotated in Encode, GeneHancer, and SuperEnhancer datasets (**Figures 1C,2**) (**Table 1**).

**Figure 2.**
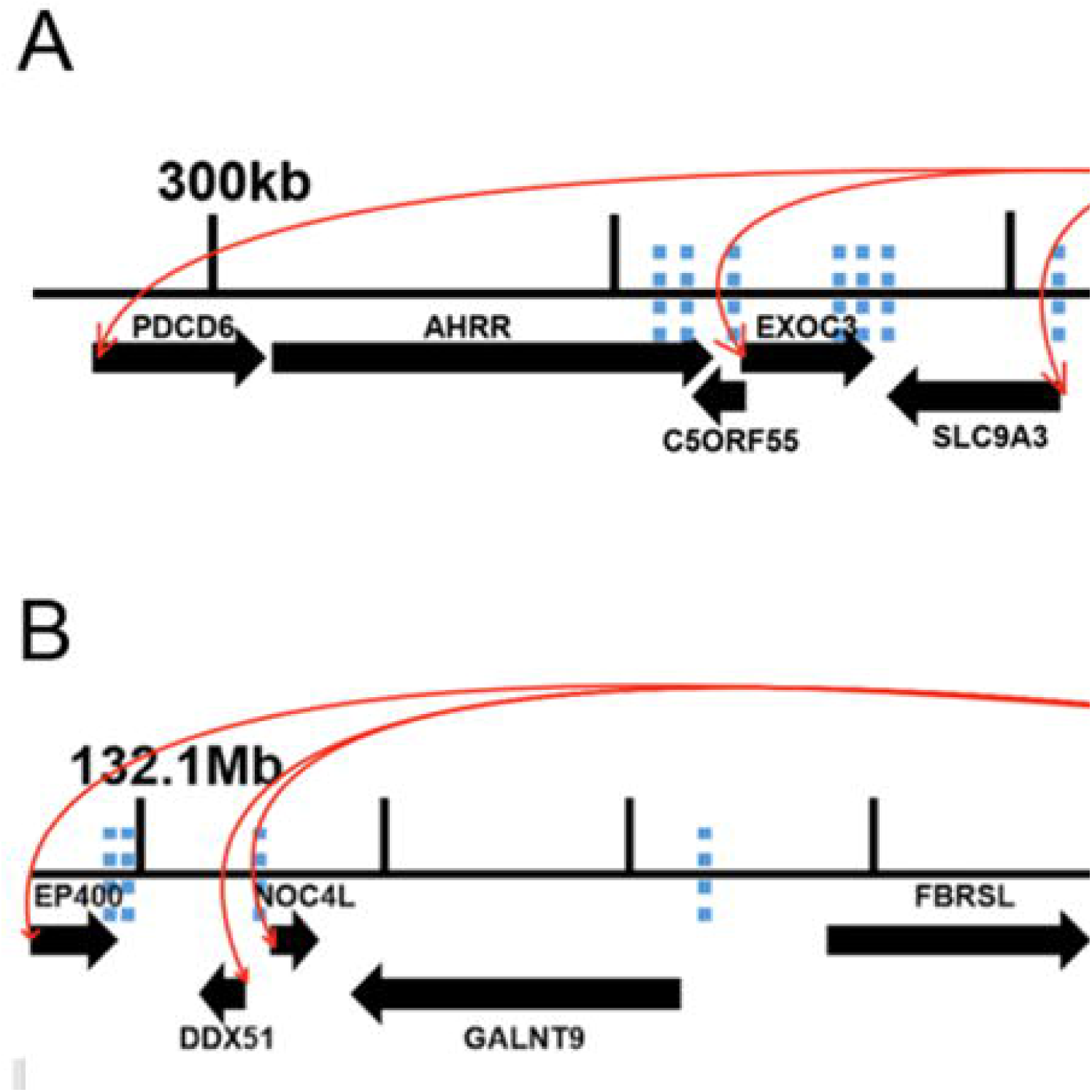
LG4 regulatory neighborhoods. Protein coding genes and their orientation as annotated in Ensembl^25^ are indicated as black arrows below the black line representing the genome. (**A**) Illustration of an insulated neighborhood within Chr5p15.33 centered on the LG4 located at GRCh38 Chr5:551935:556936:1 which is depicted above the assembly as a cartoon G4 structure labeled 550kb LG4. The red arrows originating at the LG4 and ending at various gene promoters indicate the GeneHancer annotated promoter regulations enacted by enhancers within the LG4 locus (GH05J000553, GH05J000555). The blue dashed lines represent regions of this insulated neighborhood which were concatenated to the LG4 within a single Pore-C read (Read SRR11589412.3086865.1). (**B**) Illustration of an insulated neighborhood within Chr12q24.33 which contains an LG4 located at GRCh38 Chr12:132686134:132690031:1. The LG4 is shown as a cartoon G4 structure labeled 132.68Mb LG4. The red arrows originating at the LG4 and terminating at various promoters within the assembly denote GeneHancer annotated promoter regulations imposed by the enhancer element residing within the LG4 locus (GH12J132686). The blue dashed lines denote regions of this insulated neighborhood which were joined to the LG4 within a single Pore-C read (Read SRR11589411.5760096.1). Annotations of Pore-C reads (SRR11589412.3086865.1 and SRR11589411.5760096.1) are detailed in **Supplemental Table 4.**

**Table 1.** LG4 enhancer annotations. Enhancer positions overlapping Chr5 and Chr12 LG4s and their putative gene regulations were curated from several databases including Genehancer, SEdb 2.0, Ensembl and ENCODE. LG4 positions are listed in red at the bottom of each table.

Chromosome 5 LG4 Enhancer Annotations
| Enhancer ID | Source | Start Position (Chr5) | Stop Position (Chr5) | Noted Gene Associations/Regulations |
| --- | --- | --- | --- | --- |
| GH05J000553 | GeneHancer | 553660 | 554453 | EXOC3, CEP72, BRD9, SLC9A3 |
| GH05J000555 | GeneHancer | 555169 | 557644 | PDCD6, CEP72, BRD9, SLC9A3 |
| SE_02_008600693 | SE db 2.0 | 505305 | 558774 | SLC9A3, LOC25845 |
| SE_02_024100501 | SE db 2.0 | 543481 | 558675 | SLC9A3 |
| SE_02_020700020 | SE db 2.0 | 532856 | 558643 | SLC9A3, CEP72 |
| SE_00_002000529 | SE db 2.0 | 498817 | 557908 | SLC9A3, LOC25845 |
| SE_00_003600317 | SE db 2.0 | 498742 | 557880 | SLC9A3, LOC25845 |
| SE_02_035400733 | SE db 2.0 | 473202 | 554505 | SLC9A3, LOC25845, EXOC3, C5ORF55 |
| ENSR000001256048 | Ensembl | 555201 | 555400 | Not indicated |
| ENSR000001256049 | Ensembl | 556401 | 556600 | Not indicated |
| EH38E2352151 | ENCODE | 553896 | 554206 | SLC9A3, CEP72, EXOC3 |
| EH38E2352153 | ENCODE | 555051 | 555387 | SLC9A3, CEP72, EXOC3 |
| EH38E2352157 | ENCODE | 556903 | 557118 | SLC9A3, CEP72, EXOC3 |
|  | Chr 5 LG4 | 551935 | 556936 |  |

Chromosome 12 LG4 Enhancer Annotations
| Enhancer ID | Source | Start Position (Chr12) | Stop Position (Chr12) | Noted Gene Associations/Regulations |
| --- | --- | --- | --- | --- |
| GH12J132686 | GeneHancer | 132685200 | 132690084 | POLE, PXMP2, ANKLE2, NOC4L, EP400, ZNF891, ZNF84, ZNF140, GOLGA3, CHFR, ZNF605, DDX51, PGAM5, ZNF26, ZNF10 |
| SE_02_116400065 | SE db 2.0 | 132684308 | 132695135 | PXMP2, POLE, PGAM5 |
| SE_02_055500763 | SE db 2.0 | 132685389 | 132689520 | PXMP2, POLE, PGAM5 |
| SE_02_039600318 | SE db 2.0 | 132688822 | 132789737 | ANKLE2, PXMP2, PGAM5, GOLGA3, POLE |
| SE_02_104301098 | SE db 2.0 | 132639038 | 132685737 | POLE, PXMP2, PGAM5, P2RX2, LRCOL1 |
| EH38E1657781 | ENCODE | 132686748 | 132686912 | POLE, PXMP2, ANKLE2 |
| EH38E1657782 | ENCODE | 132686991 | 132687193 | POLE, PXMP2, ANKLE2 |
| EH38E1657785 | ENCODE | 132689268 | 132689614 | POLE, PXMP2, ANKLE2 |
| EH38E1657786 | ENCODE | 132689796 | 132690144 | POLE, PXMP2, ANKLE2 |
|  | Chr 12 LG4 | 132686134 | 132690031 |  |

Next, in light of a model of G4-based enhancer:promoter interaction proposed in 2015 based on the identification of a propensity for single enhancer:promoter pairs to contain putatively interacting minimal G4 motif components^15^, we elected to examine the potential of the promoters annotated as being regulated by these LG4 enhancers to contribute to G4 formation. Analyses of the gene promoters likely regulated by LG4 enhancers find them significantly enriched with G4-capable sequences^23^ suggesting that LG4 sequences could potentially form composite G4s with distal promoters with a LG4 and interacting promoter each contributing half the sequence necessary to form a composite G4. As an example, we find the average number of G triplets occurring on either strand within 5kb upstream of the average human protein coding gene transcription start site to be ∼142. In contrast, the average number of G triplets occurring within 5kb upstream of the four genes (EXOC3, CEP72, BRD9, SLC9A3) annotated by GeneHancer^29^ as being regulated by an enhancer wholly embedded within the Chr5 LG4 (GH05J000553) is notably higher at 237 (range 184-291) (**Supplemental Table 1**). Furthermore, we find the average number of G triplets available for contributing to composite G4 formation within a single LG4 to be 74 times the number of available G triplets in the average target-gene promoter^23^ leading us to speculate that LG4 enhancers may act as long ‘Velcro-like’ regions simultaneously interacting with multiple gene promoters to coordinate their expressions (**Graphical Hypothesis**). Finally of note, a recent genomic and transcriptomic association study of 7,840 Cystic Fibrosis (CF) patients^32^ identified four single nucleotide polymorphisms (SNPs) located within the Chr5 LG4 significantly associated with CF lung disease severity (avg. p value = 2.83E-9), presumably through expression changes of EXOC3 and CEP72. This is supported by Genotype-Tissue Expression (GTEx) consortium as expression Quantitative Trait Loci (eQTL) data^33^ indicating significant correlations between these four SNPs (each of which involves a change to/from a G/C) and EXOC3 and CEP72 expressions (as well as the expressions of SLC9A3, TPPP, and ZDHHC11) (**Supplemental Table 2**).

### LG4s and putative target promoters are proximally located in the nucleus

We find several lines of evidence supporting the proposed colocalization of LG4s and their putative target promoters. (1) Chromatin conformation capture is a method used to investigate interactions between genomic loci that are not adjacent in the primary sequence. In this method, genomic DNA is first cross-linked (to preserve the spatial proximity of interacting loci) then restriction digested before being enzymatically ligated to concatenate sequences that are proximally located in the nucleus despite occurring at distinct, non-contiguous genomic positions^4^. Notably, one of these methods, Oxford Nanopore Pore-C, routinely generates concatenated reads averaging >10,000 nt in length and as such are ideally suited for revealing enhancer:promoter interactions^34^. In light of this, we elected to mine several Pore-C datasets (publically available via the NCBI SRA repository^26^) and readily identified numerous examples of individual Pore-C reads containing segments of both an LG4 enhancer and its annotated target promoters strongly supporting their spatial proximity in the nucleus (**Supplemental Tables 3,4**) (**Figure 2**). (2) The correlation between spatial proximity and frequency of chromosomal translocations at the scale of individual genes is well documented^35^. That said, we find genes potentially regulated by Chr5 and Chr12 LG4s frequently engage in local gene fusions (annotated in FusionGDB^36^) further supporting their proximity in nuclear chromatin (**Table 2**). And (3), a technique recently developed by our lab, EQuIP (Enhancer Quadruplex Immuno Precipitation) (**Figure 3A**) directly confirms enrichments for individual gene promoters in LG4-specific IPs. Notably, we find EQuIP pulldowns of the Chr5 LG4 enriched for DNA from the EXOC3 promoter (annotated as being regulated by an enhancer embedded in this LG4 (GH05J000553)^29^). Importantly, Chr5 LG4 EQuIP analyses demonstrate clear enrichments of EXOC3 promoter sequences in pulldowns using probes targeting the Chr5 LG4 enhancer whereas neither sequences^37^ (i) corresponding to the MYO10 promoter (located ∼15 Mbp downstream and not annotated as being regulated by this enhancer), nor (ii) located within the local Chr5 LG4 neighborhood but not occurring within an annotated promoter (internal AHRR) were appreciably enriched (**Figure 3B, Supplemental Figure 1**).

**Figure 3.**
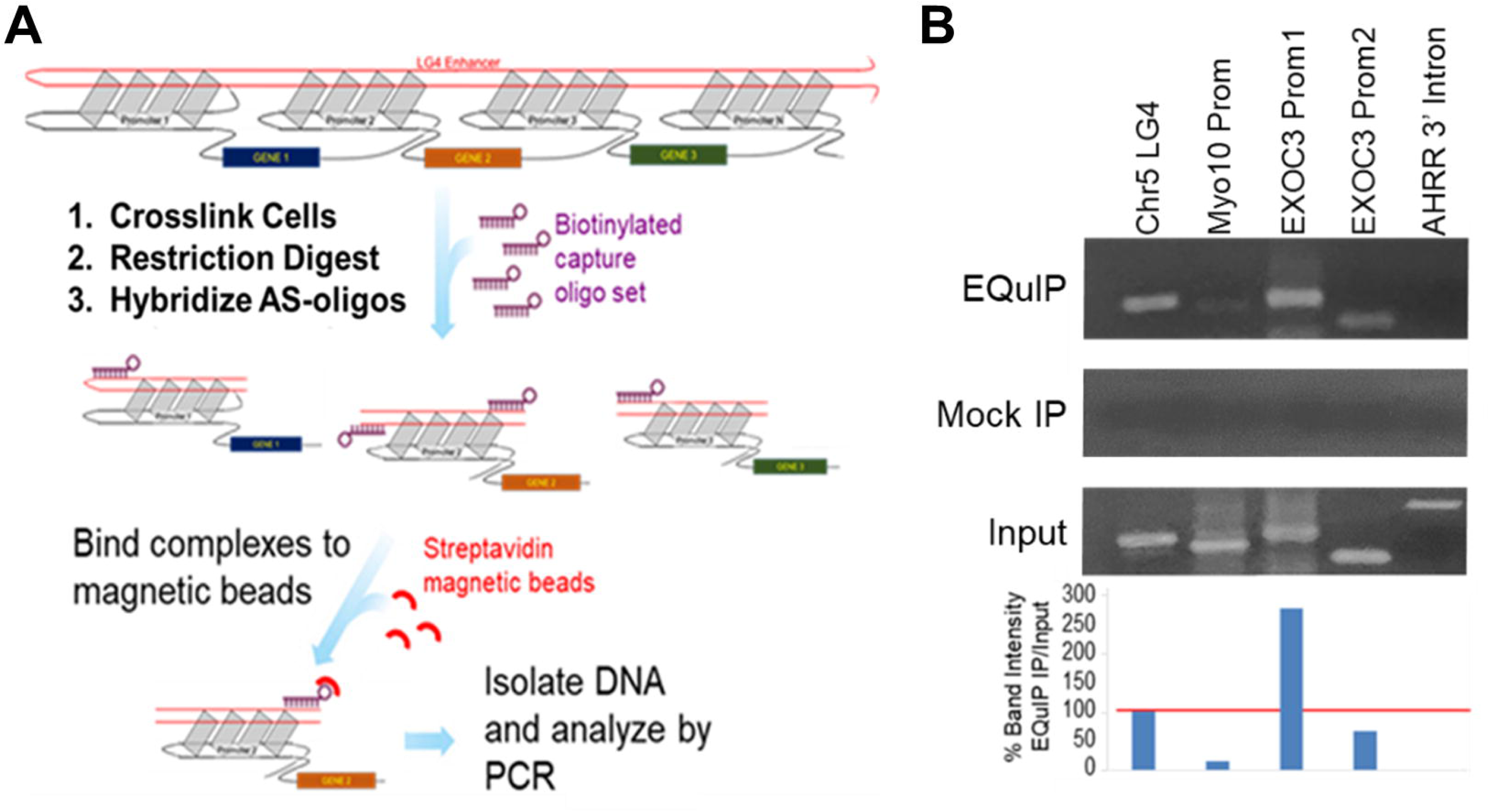
EQuIP (Enhancer Quadruplex Immuno Precipitation). (**A**) Cartoon of stepwise EQuIP protocol. In brief, EQuIP employs a probe set consisting of distinct biotinylated oligo probes complementary to different regions of the LG4. Probes are designed to target regions flanking sequences meeting the minimal criteria for G4 formation as flanking sequences are presumably held single stranded by their neighboring G4s and therefore free to basepair with complementary probes. These probe sets are combined with crosslinked, digested chromatin and allowed to hybridize to LG4 DNA. Complexes containing biotinylated-probes bound to LG4 DNA are isolated using streptavidin magnetic beads then DNA recovered and analyzed by PCR. (**B**) PCRs employing DNA template isolated from PC3 cell EQuIP pulldowns, pulldowns using nonspecific probes (Mock IP), or total input DNA collected prior to IP. PCR amplicons (∼300bp) located: (Chr5 LG4) within the Chr5 LG4, (MYO10 Prom) ∼200 bp upstream of the transcription start site of the MYO10 protein coding gene, (EXOC3 Prom1) ∼1.4 kbp upstream of the transcription start site of the EXOC3 protein coding gene, (EXOC3 Prom2) ∼2.0 kbp upstream of the transcription start site of the EXOC3 protein coding gene, (AHRR 3’ Intron) within an intron located near the 3’ end of the AHRR protein coding gene. PCR amplicons were verified by sequencing. % band intensity (EQuIP IP/Input) is presented as a bar graph below each pair of corresponding amplicons. Full gel images are shown in **Supplemental Figure 1**.

**Table 2.**
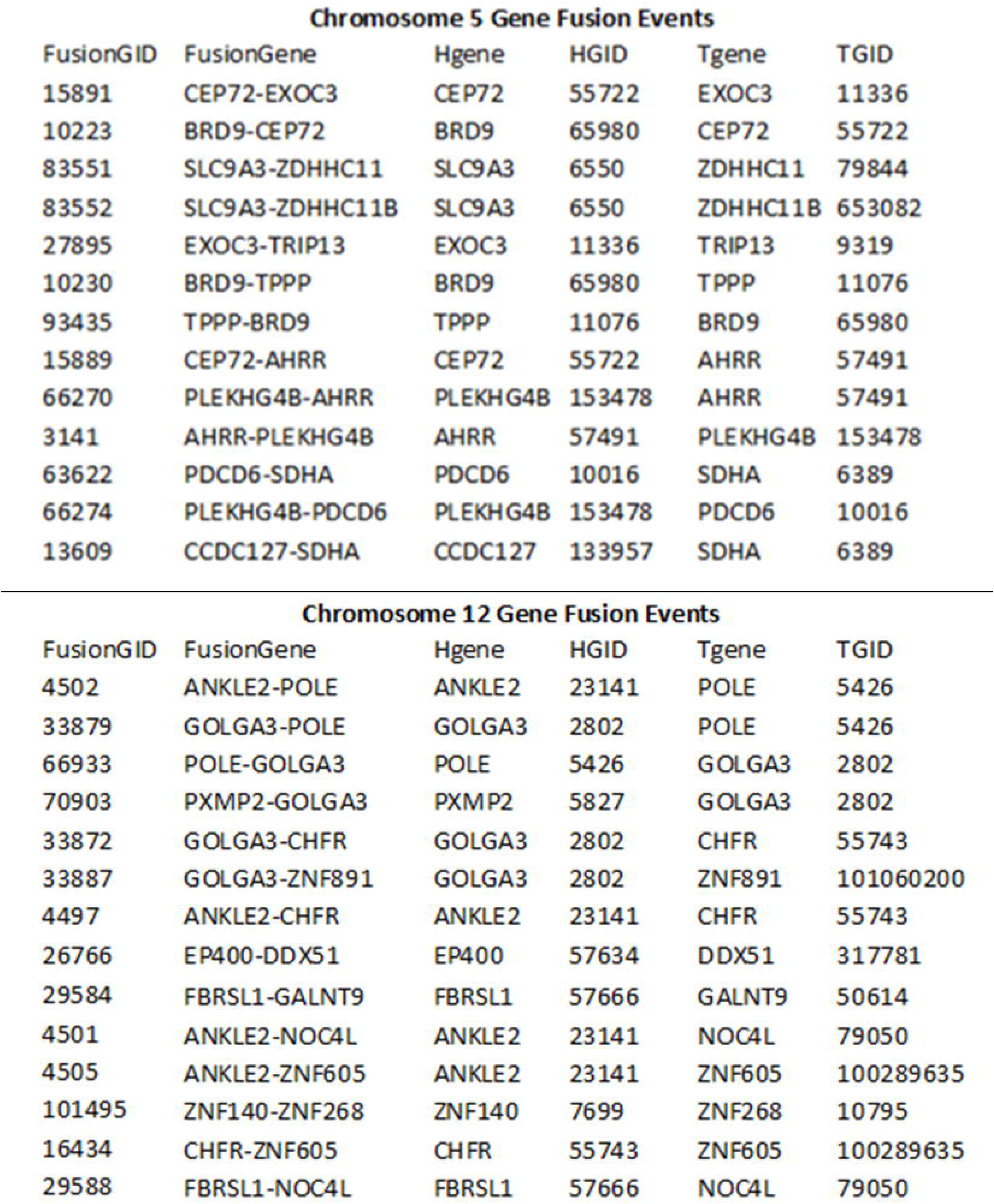
LG4 regulatory neighborhood gene fusions. Gene fusion events between genes within LG4 regulatory neighborhoods were identified using FusionGDB2.0.

### DNA sequences within the EXOC3 promoter and Chr5 LG4 can physically interact

To directly assess whether the EXOC3 promoter physically interacts with the Chr5 LG4, we performed *in-vitro* electrophoretic mobility shift assays (EMSA) designed to examine single stranded DNA (ssDNA) interactions (**Figure 4A**). Phagemids containing the EXOC3 promoter, the HIF1A promoter (a similarly sized, G4-capable promoter located on human Chr14)^38^, and the Chr5 LG4 were constructed and M13KO7 helper phage used to generate ssDNAs corresponding to each. Importantly, whereas no interaction between G4-capable HIF1A promoter and Chr5 LG4 ssDNAs was observed, nor any interaction between EXOC3 promoter and Chr5 LG4 ssDNAs after being incubated in non-G4-permissive conditions (H_2_O lacking KCl), a substantial, additive gel shift was clearly observed between ssDNAs corresponding to the + strand of the EXOC3 promoter and the - strand of the Chr5 LG4 when folded together under G4 permissive conditions (**Figure 4B**). Interestingly, a similar gel shift was also observed between ssDNAs corresponding to the - strand of the EXOC3 promoter and the + strand of the Chr5 LG4 whereas no interactions were observed between the - strand of the EXOC3 promoter and the - strand of the Chr5 LG4 nor the + strand of the EXOC3 promoter and the + strand of the Chr5 LG4 when folded together under G4 permissive conditions (**Figure 4B**). Of note, each of our phagemid generated ssDNAs migrates markedly faster after being incubated in G4 permissive (+KCl) versus non-permissive (H_2_0) conditions, regardless of their ability to independently form G4 DNA (**Supplemental Figure 2**), indicating KCl concentration alone can alter gel mobility (consistent with previous reports^39^). That said, as the size of the observed gel shift closely approximates the predicted size of an intermolecular interaction between EXOC3 promoter and Chr5 LG4 ssDNAs (**Supplemental Figure 2**), these results strongly suggest that DNA sequences within the EXOC3 promoter and Chr5 LG4 can directly, physically interact with one another and that G4 formation is required for this interaction to occur. Importantly, whereas we find EXOC3 and HIF1A promoter ssDNAs similarly capable of independently forming G4 structures, in contrast to EXOC3 promoter ssDNA, we find no evidence of an interaction between HIF1A promoter and Chr5 LG4 ssDNAs folded together under G4 permissive conditions (**Figure 4C**). We suggest this likely indicates that (i) while G4 formation is necessary for G4-mediated enhancer:promoter interactions, it is not sufficient, and (ii) additional sequence complementarities (potentially involved with mediating enhancer:promoter interaction specificity) are likely required. Finally, in an attempt to better define the sequences involved with mediating the observed EXOC3:Chr5 LG4 interactions, we elected to generate a series of truncated EXOC3 promoter constructs (**Figure 5A**). Notably, we find a 982 bp portion of the EXOC3 promoter located between 2,108 and 1,126 bp upstream of the primary EXOC3 transcription start site (TSS) is independently capable of interacting with the Chr5 LG4. Interestingly, although several minimal G4-capable sequences are located between the TSS and this region (TSS to 1,126 upstream), we find the 982 bp mediating the interaction with the Chr5 LG4 entirely devoid of any traditional G4-capable motifs (**Supplemental Figure 3**). As this clearly argues against interactions between G4s independently formed in both of the contributing sequences, we suggest these results instead agree with the general model proposed in our **Graphical Hypothesis** in which LG4 enhancers physically interact with gene promoters by forming composite G4 structures where both the LG4 and cognate promoter contribute half of the necessary sequence for G4 formation.

**Figure 4.**
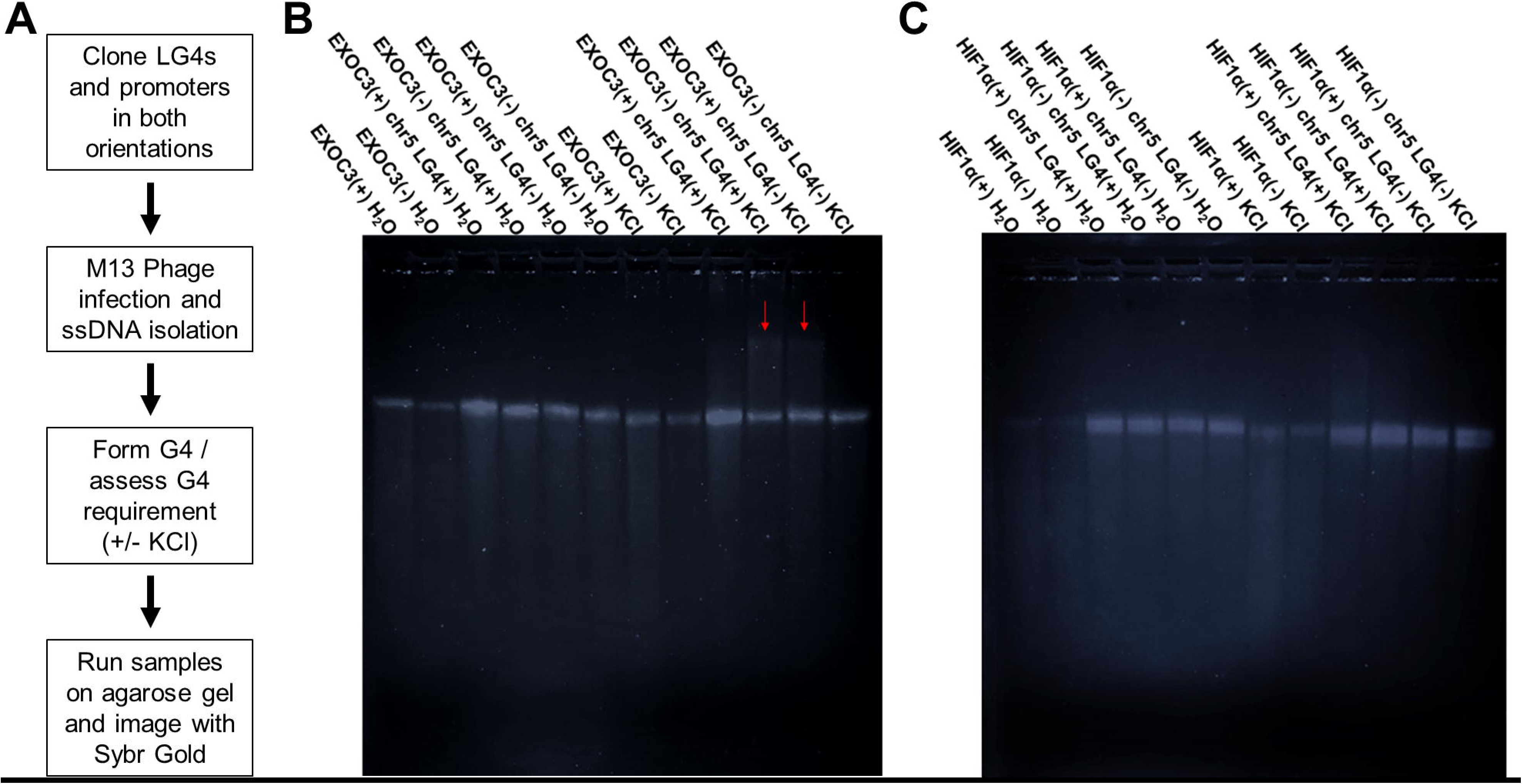
Direct interaction between EXOC3 promoter and Chr5 LG4 enhancer. (**A**) Flow chart of EMSA protocol. (**B**) Image of 1.5% agarose Tris-glycine gel ran at 4°C for 8 h at 75 V and then stained for 24 h with SYBR GOLD. Each construct was run on the gel in either the unfolded (H2O) or folded (KCl) state as indicated above the image. Samples including more than one construct were folded together. Red arrow denotes gel shift observed when the EXOC3 promoter and Chr5 LG4 are folded together. (**C**) EMSA identically performed as in **B** except the HIF1A promoter was used in place of the EXOC3 promoter.

**Figure 5.**
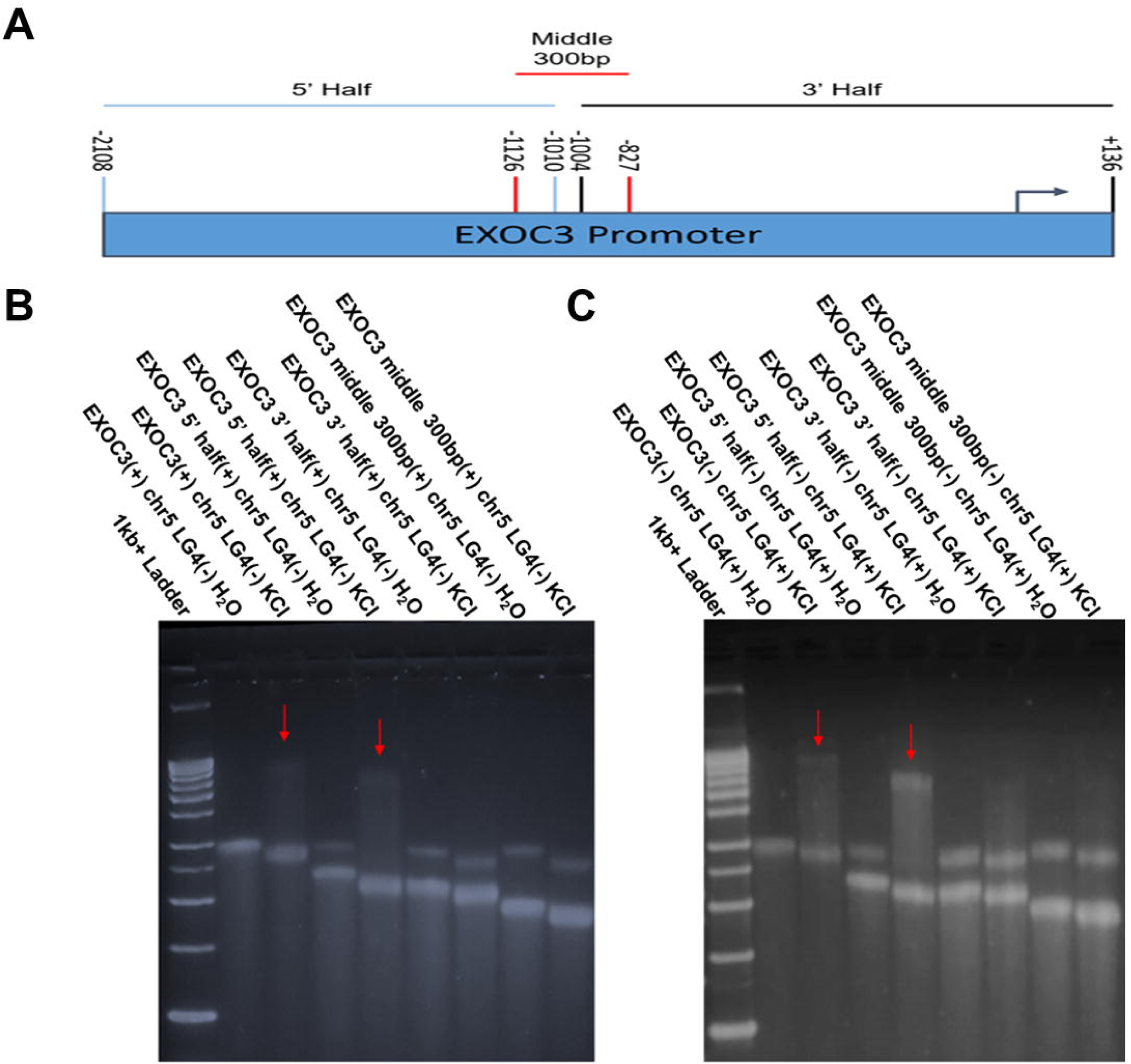
EXOC3 deletion analysis. (**A**) Cartoon depiction of positions of deletion constructs within the full length EXOC3 promoter sequence used in Figure 4 EMSA. Nucleotide positions are given with respect to the transcription start site (TSS) indicated by an arrow. (**B,C**) Images of 1.5% agarose Tris-glycine gels ran at 4°C for 8 h at 75 V then stained with SYBR GOLD for 24 h. Sample content within each well is indicated above the gel image. The strand of each construct used in this EMSA is denoted as either (+) for the sense strand or (-) for the antisense strand.

### DNA sequences within the GOLGA3 promoter and Chr12 LG4 can physically interact

To determine whether the ability of EXOC3 promoter and Chr5 LG4 ssDNAs to directly interact was unique to this locus or instead if other LG4 enhancers might similarly physically interact with their cognate promoters through a direct G4:G4 DNA based mechanism, we elected to examine a second unrelated LG4 overlapping annotated enhancers on Chr12 (**Figure 2B**). Excitingly, as with the Chr5 LG4, after being incubated in G4-permissive conditions (+KCl), a substantial, additive gel shift was clearly observed between ssDNAs corresponding to the + strand of the GOLGA3 promoter and the - strand of the Chr12 LG4 when folded together under G4 permissive conditions (**Figure 6**). The GOLGA3:Chr12 LG4 interaction proved even more specific than the EXOC3:Chr5 LG4 interaction, as no other interactions were observed between ssDNAs corresponding to the - strand of the GOLGA3 promoter and the + strand of the Chr12 LG4, the - strand of the GOLGA3 promoter and the - strand of the Chr12 LG4, nor the + strand of the GOLGA3 promoter and the + strand of the Chr12 LG4 when folded together under G4 permissive conditions (**Figure 6**). Finally, as with the EXOC3:Chr5 LG4 interaction, we find that G4 formation is likewise required for the GOLGA3:Chr12 LG4 interaction to occur and that the size of the gel shift observed closely approximates the predicted size of an intermolecular interaction between these two ssDNAs (**Supplemental Figure 2**) demonstrating that DNA sequences within the GOLGA3 promoter and Chr12 LG4 can also directly, physically interact with one another.

**Figure 6.**
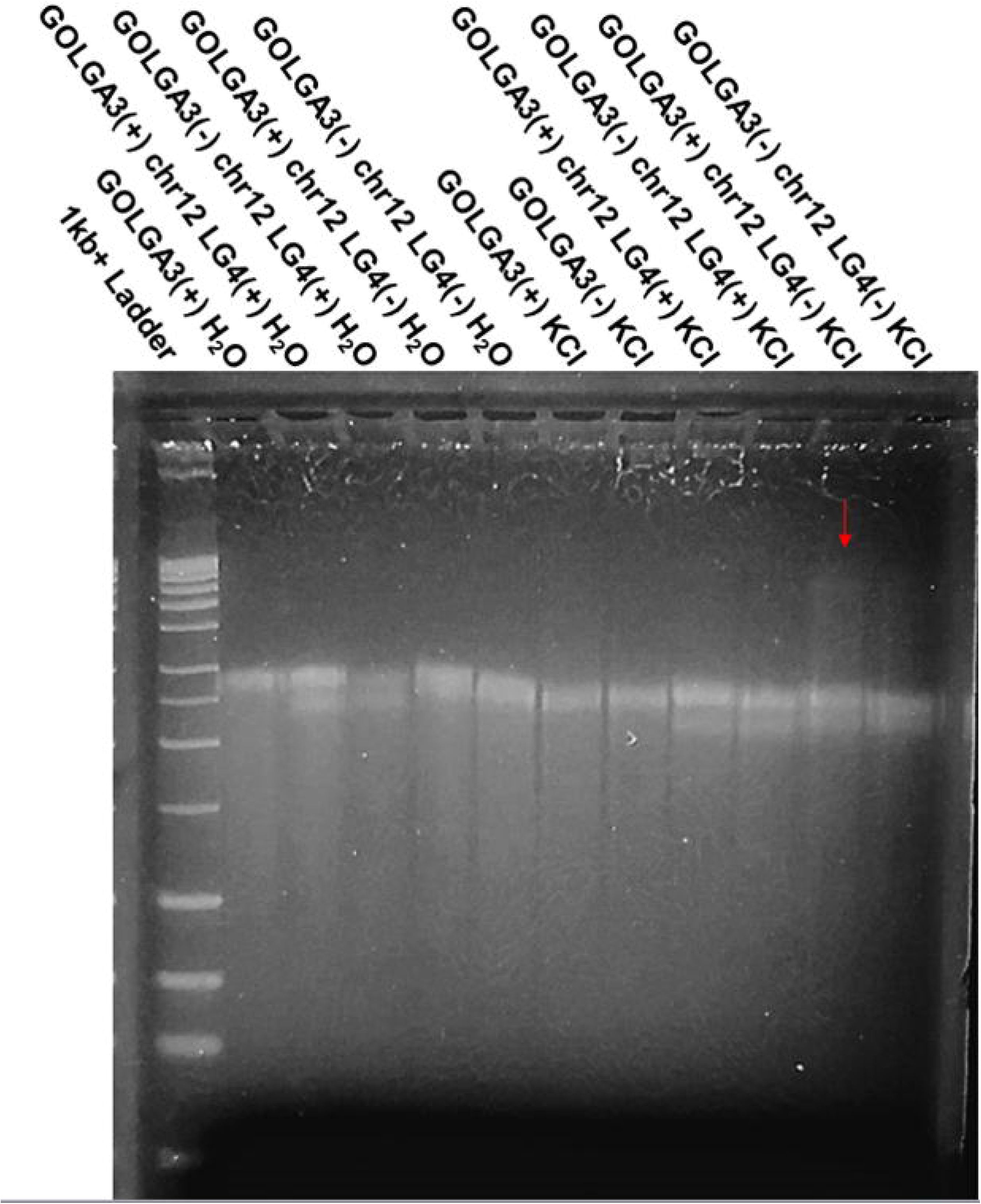
Direct interaction between GOLGA3 promoter and Chr12 LG4 enhancer. Image of 1.5% agarose Tris-glycine gel ran at 4°C for 8 h at 75 V and then stained for 24 h with SYBR GOLD. Each construct was run on the gel in either the unfolded (H2O) or folded (KCl) state and is indicated above the image. Samples including more than one construct were folded together. The red arrow denotes the gel shift observed when the GOLGA3 promoter and Chr12 LG4 are folded together.

## Discussion

Long G4-capable (LG4) loci are defined as genomic regions densely populated with minimal G4 capable sequences (GGGnGGGnGGGnGGG). Our previous work demonstrated that LG4s can assume higher order structures^23^ although the requirements for and determinants of their assembly have not been fully elucidated. Moreover, although 217 of 301 previously identified LG4s overlapping annotated enhancer elements, no direct participation of G4 structures within LG4s with regard to gene regulation have been reported to date. That said, despite the current generally accepted model for enhancer:promoter communication involving enhancer:promoter engagement being facilitated by interactions between activator proteins independently bound to each^1,11^, this study was designed to test an alternative hypothesis: that LG4 enhancers physically interact with their cognate promoters via a direct G4:G4 DNA based mechanism.

Notably, numerous studies have suggested potential roles for G4 DNA in enhancer activity to date. As examples(i) G4 ChIP-Seq has repeatedly shown G4 structures are enriched in promoters and enhancers, (ii) G4 formation is significantly correlated with elevated transcriptional activity^40,41^, and (iii) a genome-wide mapping of G4 structures in 2022 found G4s located in enhancers are preferentially formed by inter-strand G4 folding (e.g., composite G4s containing only two GGG repeats from one strand of an enhancer)^42^. That said, a model for G4-based enhancer:promoter interaction was first proposed in 2015 based on the identification of a marked propensity for single enhancer:promoter pairs to contain potentially interacting minimal G4 motif components^15^. In 2019, Hou et al. proposed a similar model for G4-based enhancer:promoter interactions after finding more than 99% of G4s overlap known transcription factor binding sites (TFBSs), G4s are significantly enriched at boundaries of topological associated domains (TADs), and that frequent interactions occur between G4-containing regulatory elements^43^. While it is clear that enhancers regulate distal genes by genomic looping and physical interaction, how enhancers target the right genes remains unclear. The current paradigm for how enhancer:promoter specificity is primarily mediated involves a model in which the genome is divided into a hierarchical series of domains with enhancers regulating target promoters only within their own individual domains^44^. At the top level, chromosomes occupy distinct spaces in the nucleus deemed chromosome territories^45^ which can be divided into ∼1 Mbp regions known as TADs^46^. TADs are themselves comprised of a series of smaller DNA loops known as insulated neighborhoods (INs) formed by interactions between two DNA sites bound by CTCF and the cohesion complex and typically contain 1 to 8 genes and 1 or 2 enhancers^47^ (similar to the regulatory neighborhoods proposed in **Figure 2**). Enhancers located within TADs are believed to only interact with genes located in their respective neighborhood^48^ as when TAD boundaries are disrupted, aberrant enhancer:promoter interactions result in significant gene misexpressions^49–51^. Notably, Hou et al. found that G4s are significantly enriched at TAD boundaries and that G4 content strongly correlates with occupancy of architectural proteins critical for TAD formation. What’s more, they found that adjacent, G4-containing boundaries frequently interact, the insulation abilities of CTCF binding sites and TAD boundaries are significantly reinforced by G4s, more than 99% of G4s at these positions overlap TFBSs, and that CTCF and cohesin binding sites are preferentially located near G4s. Collectively, these findings clearly support G4 involvement in loop extrusion and distal interactions between enhancers and promoters^43^. We similarly previously identified, significant associations (p < 0.0001) between our LG4 loci and ChIP-seq peaks corresponding to 26 different TFs including CTCF, YY1, and SP1^23^ (known to associate with promoter G4s^52^). Although G4 DNA was only recently directly linked to CTCF recruitment^53^, roles for CTCF and cohesin in 3D genomic organization^54^ and for CTCF promoter binding as a mediator of long distance enhancer dependent transcription^28^ have long been established. YY1 has also been shown to contribute to enhancer:promoter interactions^55^, and in 2021, Li et al. identified G4 binding as a molecular requirement for YY1-driven long range enhancer:promoter interactions^56^. Notably, they found displacement of YY1 from G4 structures with small-molecule G4 ligands, unwinding of G4 structures by overexpression of BLM helicase, and CRISPR–Cas9 mutation of G4-forming promoter sequences could each disrupt YY1-mediated DNA looping. In addition, they also showed expression of genes harboring G4 structures in their promoters could be significantly perturbed by either inhibiting YY1:G4 binding (by the use of G4-stabilizing ligands PDS or TMPyP4) or RNAi depletion of YY1^56^. Finally of note, in 2023 Roy et al. artificially inserted an array of G4s taken from a known enhancer into a genomic locus devoid of any G4 rich regions^57^. Subsequent Hi-C experiments found the introduced G4s engaged in several long-range interactions and led to increased gene expressions up to 5 Mb away, local enrichment of H3K4Me1 and H3K27Ac enhancer signals, and recruitment of transcriptional coactivators directly supporting roles for G4s as enhancers and in mediating long-range chromatin interactions^57^.

That said, despite a mounting body of evidence implicating G4 involvement, the current, generally accepted model for enhancer:promoter communication posits that enhancer:promoter engagement is dictated by interactions between activator proteins independently bound to each^1,11^. The work detailed in this report, however, supports a direct role for enhancer and promoter DNA sequence interaction in mediating enhancer:promoter communication and specificity. Importantly, we find several lines of evidence indicating LG4s and their putative target promoters are proximally located in the nucleus. (1) We identify numerous individual chromatin conformation capture reads containing concatenated enhancer and promoter sequences (**Supplemental Table 3**). (2) We find genes potentially regulated by Chr5 and Chr12 LG4s frequently engage in local gene fusions (annotated in FusionGDB^36^) **(Table 2**). And (3) EQuIP pulldowns using probes specific for Chr5 LG4 DNA are highly enriched with DNA from the EXOC3 promoter (**Figure 3**). In addition to simply establishing that LG4s and their putative target promoters are proximally located, our work also experimentally confirms direct, specific interactions between ssDNAs corresponding to (1) a LG4 found on human Chromosome 5 (Chr5 LG4) and the EXOC3 promoter, one of its predicted target promoters located just over 100 kbp upstream (**Figure 4**) and (2) a LG4 found on human Chromosome 12 (Chr12 LG4) and the GOLGA3 promoter, one of its predicted target promoters located nearly 200 kbp downstream (**Figure 6**). Notably, these promoter:LG4 interactions were only detected when participating ssDNAs were folded together under G4 permissive conditions (+ KCl) and not in the absence of a stabilizing cation (H_2_O) indicating that G4 formation is required for these DNA-based enhancer:promoter interactions. Furthermore, we find a 982 bp portion of the EXOC3 promoter is required for and independently capable of interacting with the Chr5 LG4 (**Figure 5**), and interestingly, that this 982 bp portion is entirely devoid of any minimal G4 capable sequence motifs (**Supplemental Figure 3**). As such, we suggest the apparent inability of this region to form independent G4 structures coupled with the observation that G4 formation is required for this region to engage in DNA-based enhancer:promoter interaction is clearly in agreement with the general model proposed in the **Graphical Hypothesis** in which LG4 enhancers physically interact with gene promoters by forming composite G4 structures where both the LG4 and cognate promoter contribute half of the necessary sequence for G4 formation.

Additionally of note, the data presented in this study also suggests that individual LG4 enhancers regulate a specific set of target promoters and potentially coordinates their expressions. We find the average number of G triplets (GGG) potentially contributing to composite G4 formation within a single LG4 is 74 times the number of available G triplets found in the average inferred target-gene promoter^23^ leading us to speculate that the high number of available G4 donor sequences within a single LG4 may allow LG4 enhancers to act as long “Velcro-like” regions that simultaneously interact with a number of neighboring gene promoters coordinating their expressions (see **Graphical Hypothesis**). Although conclusively establishing a role for LG4s in promoter coordination is beyond the scope of the current study and will ultimately require more extensive examination, we find several lines of evidence clearly suggesting that the expressions of genes regulated by LG4 enhancers are likely coordinated (**Supplemental Figure 4**). Particularly of note, our preliminary assessment of TCGA patient expression data^58^ finds the expressions of CEP72, BRD9, and PDCD6 to be significantly, positively correlated with EXOC3 expression in lung adenocarcinoma, and similarly, the expressions of EP400, ZNF26, ZNF84, and ZNF605 to be significantly, positively correlated with GOLGA3 expression in sarcoma.

Regardless of their ability to coordinate gene expressions, the regulatory action which LG4s impose on their respective target genes is likely disease relevant. Particularly of note to this report, in 2020, a GWAS designed to identify genomic modifiers of Cystic fibrosis (CF) identified 28 genes (near known GWAS loci) whose expressions are highly associated with CF lung disease severity in patients^59^. Strikingly, of these 28 disease modifying genes, 12 (43%) (e.g., BRD9, CEP72, EXOC3, TPPP, ZDHHC11) reside in the Chr5 LG4 neighborhood depicted in **Figure 2A**, and what’s more, the expressions of CEP72, EXOC3, TPPP and ZDHHC11 were identified as the four most highly associated with CF severity genome wide.

Additionally of note, a significant number of SNPs identified as associated with EXOC3, CEP72, and TPPP expressions were not located within the respective promoters of these genes but instead the strongest signals were within an ∼40kb window centered around the Chr5 LG4^59^. Most recently, a genomic and transcriptomic association study of 7,840 Cystic Fibrosis (CF) patients^32^ identified four CF-relevant SNPs (located within the Chr5 LG4) significantly associated with the expressions of EXOC3 and CEP72 (**Supplemental Table 2**). Collectively these findings suggest that perturbation of the central LG4 enhancer regulating these genes may have substantive consequences on CF lung disease phenotype. More directly, these SNPs showed significant eQTLs with the expression of these genes by GTEx in multiple human tissues including lung^33,59–64^. Although we have not yet attempted to comprehensively identify associations between any of the other 300 LG4s identified in our original study and any other diseases / genetic networks, it is tempting to speculate that regulatory networks associated with additional LG4s may prove similarly relevant to other diseases.

Other topics not fully addressed in the current study but clearly warranting further examination include determining how promoter:LG4 interactions are regulated/function in vivo, fully defining the sequence requirements mediating interaction and promoter specificity, and determining how widespread G4-based enhancer:promoter interactions actually are. Although the work detailed here biochemically establishes direct interaction between LG4 enhancers and their cognate promoters, a major limitation of the current study is that it does not explore the cellular context required for these interactions occur. It is entirely possible that a particular LG4 enhancer may regulate one set of promoters in one cell type and regulate another group of promoters or to be inactive altogether in other cell types. Also of note, the current study was restricted to an examination of just two distinct LG4s limiting our ability to confidently conclude that the regulatory mechanism described in this work represents a general mechanism of LG4 action. Furthermore, while our work successfully (1) provided multiple lines of evidence indicating that specific LG4s and their target promoters are proximally located in the nucleus, and (2) biochemically demonstrated that ssDNAs corresponding to individual LG4s and promoters specifically associate with each other in a G4-dependent manner, establishing clear roles for G4-mediated targeting in facilitating enhancer:promoter specificity in vivo will clearly require additional study.

In summary, in contrast to the generally accepted model for enhancer:promoter communication in which enhancer:promoter engagement is dictated by interactions between activator proteins, this work describes a novel G4 DNA-based mechanism capable of mediating enhancer:promoter interaction in vitro and confirms LG4s and their target promoters are proximally located in the nucleus supporting the likely relevance of this mechanism in vivo. In addition, we also find evidence indicating that perturbation of a LG4 enhancer located on human chromosome 5 may have direct consequences on CF lung disease severity potentially suggesting that additional LG4s may be similarly associated with other diseases and that G4 stabilizing/destabilizing therapies may prove more widely applicable than currently appreciated.

## Methods

### Detailing LG4 enhancer overlaps, promoter G4 capacity, and LG4 loci gene fusions

LG4 and gene promoter locations were taken as occurring in Human Genome Release 77 (hg38)^25^. Potential enhancer regulations were obtained from: (1) Ensembl Regulatory Build^24^, (2) a comprehensive Super-Enhancer database (SEdb 2.0)^65^, (3) UCSC genome browser Encode annotations^30^, and (4) the GeneHancer DB^29^. Significant enrichment of potential G4 contributing sequences (e.g., GGG) in LG4 neighborhood promoters (or randomly selected, size matched promoters not proximal to an LG4) was calculated using chi-square after individual motifs (e.g., GGG) were enumerated using an in house python script. Full FusionGDB gene fusion datasets^36^ were downloaded and fusions between genes in LG4 neighborhoods (or randomly selected, size matched control locations) identified then significance calculated using an unpaired two-tailed t-test (as in ^23^).

### Analysis of Pore-C datasets

First, all >100 bp sequences perfectly mapping to multiple places in the human genome (hg38)^25^ occurring within EXOC3 and GOLGA3 promoters were masked and excluded from consideration. Resulting masked promoter sequences were then aligned to individual reads obtained from publicly available pore-C datasets housed in the NCBI SRA database^26^ via BLAST^66^. BLAST parameters were set at 100 max target sequences, word size 15, and an expected alignment threshold of e value= 1×10^15^. Match/mismatch scoring parameters were 2,-3. Gap costs were existence: 5 and extension: 2 and alignments to low complexity regions were allowed. The 100 highest scoring pore-C reads aligning to each promoter in individual datasets were identified then aligned to the human genome using the blastn tool in the Ensembl genome browser^25^ with search parameters set at 500 max target sequences, word size 11, and an expected alignment threshold of e value= 1×10^-50^. Scoring parameters consisted of match/mismatch scores (1,-3), and gap penalties enforced as opening: 2, extension 2. Alignments to low complexity regions were allowed and query sequences were not filtered using RepeatMasker. In the event that a portion of a pore-C read aligned to more than one place, the alignment with the lowest e-value was considered the bonafide alignment. Individual reads containing sections of both the Chr5LG4 and EXOC3 promoter or Chr12LG4 and GOLGA3 promoter were identified then the origin of each fragment contained within a given read similarly defined by genomic alignment (**Supplemental Tables 3, 4)**.

### Enhancer Quadruplex Immunoprecipitation (EQuIP)

PC3 cells were grown to 100% confluence in a T75 flask then crosslinked with glutaraldehyde (Fisher Scientific Hampton, NH cat no. O2957-1) diluted to 1% in PBS (Gibco Waltham, MA cat no. 10010049). Crosslinking reactions were incubated for 10 min at room temperature (RT) while slowly mixing on an orbital shaker. Crosslinking was then quenched by adding 1 mL of 10x glycine (Thermo Fisher Scientific Waltham, MA cat no. 043497-36) then incubating 5 min at RT followed by 10 min on ice. At this point, crosslinked cells were either permeabilized for downstream procedures or flash frozen and stored at - 80°C for future use. To permeabilize cells, they were resuspended in a mix of 500 µL of 4°C permeabilization buffer (10 mM Tris-HCl pH 8.0 (Fisher Scientific cat no. 77-86-1), 10 mM NaCl (Fisher Scientific cat no. S25541A), 0.2% IGEPAL CA-630 (Sigma-Aldrich St. Louis, MO cat no. I3021) supplemented with 50 µL protease inhibitor cocktail III (Thermo Fisher Scientific cat no. J64283-LQ) and incubating on ice for 15 min. Cells were then pelleted by centrifugation at 500 x g for 10 min and washed with 200 µL of 1.5X DpnII reaction buffer (New England Biolabs Ipswich, MA cat no. R0543S), pelleted again and then resuspended in 300 µl of cold 1.5X DpnII reaction buffer (NEB cat no. R0543S). Chromatin were denatured by adding 33.5 μL of 1% SDS (Fisher Scientific cat no. BP166-100) then incubated for exactly 10 min at 65°C with gentle agitation then placed immediately on ice. SDS was quenched by adding 37.5 μL of 10% Triton X-100 (Sigma-Aldrich cat no. 93443) then incubating on ice for 10 min. Cells were pelleted briefly (pulsed at 500 x g) to remove supernatant, then digested with a final concentration of 1 U/μL of DpnII (NEB cat no. R0543S) in 450 μL of 1X digestion reaction buffer. Reactions were gently mixed by inversion and incubated at 37°C for 18 h while shaking at 250 rpm to prevent condensation. The restriction digest was then heat inactivated by incubating at 65°C for 20 min. Cells were then pelleted by centrifugation at 500 x g for 1 min to remove supernatant and then resuspended in a complete T4 ligation reaction mix consisting of 100 µL 10X T4 ligase buffer, 10 µL 10 mg/mL BSA (NEB cat no. B9000s), 840 μL DNA grade H_2_O (Invitrogen Waltham, MA cat no. AM9922), and 50 μL T4 DNA Ligase (NEB cat no.M0202L). Ligation reactions were cooled to 16°C and incubated for 6 h. Upon completion of the ligation reaction, cells were reverse crosslinked by treatment with 100 μL (20 mg/mL) Proteinase K (Thermo Fisher Scientific cat no. FEREo0491), 100 μL 10% SDS (Fisher Scientific cat no. BP166-100) and 500 μL 20% Tween-20 (Fisher Scientific cat no. BP337-500) in a total volume of 2000 μL with nuclease free water (Invitrogen cat no. AM9922) and incubated at 56°C for 18 h. After reverse crosslinking, samples were pelleted by centrifugation at 16,000 x g for 1 min and resuspended in complete lysis buffer consisting of 20 μL protease cocktail inhibitor III (Thermo Scientific cat no. J64283-LQ), 5 μL RNase A (Invitrogen cat no. 46-7604), 1000 μL lysis buffer (50 mM Tris-HCl pH 7.0 (Fisher Scientific cat no. 77-86-1) 10 mM EDTA (Fisher Scientific cat no. S311-500), 1% SDS (Fisher Scientific cat no. BP166-100)). If no pellet is observed after centrifugation of reverse-crosslinking reaction, then proceed with hybridization using the reverse crosslinking reaction mix in place of lysis buffer. 1.7 mL Hybridization buffer (750 mM NaCl (Fisher Scientific cat no. S25541A), 50 mM Tris-HCl pH 7.0 (Fisher Scientific cat no. 77-86-1), 1 mM EDTA (Fisher Scientific cat no. S311-500), 1% SDS (Fisher Scientific cat no. BP166-100)) supplemented with 300 μL formamide (Fisher Scientific cat no. BP227-500), and 10 μL RNase A (Invitrogen cat no. 46-7604) was added for every 1 mL of cell lysate (or reverse crosslinking reaction). 100 pM of Biotinylated probes (Integrated DNA Technologies Coralville, IA) antisense to the Chr5LG4 (primer probe table) or non-targeting control probes (or no probes for input DNA control) were added to the hybridization mix and incubated at 37°C for 4 h in an orbital shaker (250 rpm) following this step the sample designated for input DNA was isolated by phenol-chloroform isoamyl alcohol 25:24:1 (Invitrogen cat no. 15593-031) extraction. To perform streptavidin pulldown, 120 μL (4 mg/mL) streptavidin magnetic beads (NEB cat no. 50-812-660) were washed 3 times with 1 mL of lysis buffer then resuspended in 100 μL of complete lysis buffer and added to the hybridization mix and then incubated at 37 °C for an additional 30 min with mixing. Tubes were placed on a magnetic separator until solution was clear. Supernatant was aspirated and streptavidin magnetic beads were washed once with 1 mL of pre-warmed wash buffer consisting of 2X SSC (Thermo Fisher Scientific cat no. AM9763), 0.5% SDS (Fisher Scientific cat no. BP166-100), 1 mM AEBSF (Fisher Scientific cat no. AC328110500) supplemented with 20 μL protease cocktail inhibitor III (Thermo Scientific cat no. J64283-LQ). Tubes were placed back on magnet until solution became clear, the supernatant was discarded and the DNA was eluted from the magnetic beads by adding 150 μL of DNA grade H_2_O (Invitrogen cat no. AM9922) and incubating at 37°C for 30 min while shaking (250 rpm). To maximize yield the elution step can be repeated a second time. DNA was quantified by Qubit (Thermo Fisher scientific cat no. Q33238) and used as template DNA for PCR to verify genomic interactions with the Chr5 LG4. If eluted DNA is insufficiently pure, a subsequent extraction by phenol-chloroform isoamyl alcohol 25:24:1 can be performed. PCRs were performed using primers specific to regions suspected to interact with the Chr5 LG4 or to control regions not predicted to interact with the Chr5 LG4. The primer sequences used in each reaction can be found in the primer probe table. PCRs were performed in 50 μL reactions with 1 μL (10 μM) forward primer, 1 μL (10 μM) reverse primer, 3 μL MgCl_2_ (25 mM), 2 μL DNTP (10 mM each), 5 μL 10X Taq buffer, 1.25 μL Taq polymerase (1 U/μL) (Fisher Scientific cat no. FEREP0404), and 61 pg of EQuIP DNA or 10 ng input DNA filled to a 50 μL with DNA grade H_2_O (Invitrogen cat no. AM9922). The thermal cycling parameters for these reactions were an initial denaturation at 95°C for 1 min, followed by 36 cycles of 30 sec at 95°C, 30 sec at 57°C, 25 sec at 72°C, then a final extension step at 72°C for 1 min. PCR products were then run on a 1% agarose TBE gel for 50 min at 100 V and stained with EtBr (Fisher Scientific cat no. BP102-5).

### Electrophoretic Mobility Shift Assay (EMSA)

All LG4 and promoter elements used in electrophoretic mobility shift assays were cloned into TOPO TA pCR2.1 (Invitrogen cat no. 45-064-1) in both the sense and antisense orientations and verified by sequencing. Confirmed constructs were transformed into F’*Iq* Competent *E. coli* (NEB cat no. C2992H) and resulting bacterial colonies used to inoculate 50 mL of LB and grown at 37°C for 6 h at 250 rpm in an orbital shaker. Next, M13KO7 helper phage (New England Biolabs N0315S) was added to each culture (final concentration of 1 x 10^8^ pfu/mL) and incubated at 37°C for 1.5 h at 250 rpm after which kanamycin was added (final concentration 70 µg/ml) then grown overnight at 37°C at 250 rpm. ssDNA was isolated the following day per M13KO7 helper phage standard manufacturer protocol (NEB cat no. N0315S).

ssDNA constructs generated by M13KO7 helper phage were run on a 1% agarose 1X TBE gel at 100 V for 45 min then size selected ssDNA purified by gel extraction using Wizard SV Gel and PCR Clean-Up System (Promega Madison, WI cat no. PR-A9281) and quantified via Nanodrop 6000 (Thermo-Scientific). 20 µl (15 ng/µl) of each ssDNA promoter were boiled either separately or together with 10 μL (10 ng/μL) of LG4 ssDNA (for samples containing only LG4, 20 μL of LG4 ssDNA was used) at 98°C for 10 min then held at 80°C for 10 min during which pre-heated KCl solution (to fold G4) (final conc. 250 mM) or an equal volume of pre-heated ultrapure H_2_O (unfolded controls) was added to the indicated samples and slow cooled to 45°C over 1 h then held at 16°C. Resulting ssDNAs folded in either KCl or water were next ran on a 1.5% agarose 1X Tris-glycine (Bio-Rad Hercules, CA cat no. 1610734) gel at 75 V for 8 h at 4°C after which gels were stained for 24 h with SYBR GOLD (Invitrogen cat. S11494) diluted 1:10000 in 1X Tris-glycine (Bio-Rad cat no. 1610734) then imaged on a UV Transilluminator FBTIV-88 (Fisher Scientific).

### Plasmid Construction

*Topo TA pCR2.1 plasmids*. The EXOC3 promoter, GOLGA3 promoter, HIF1A promoter, Chr5LG4 and Chr12 LG4 were each PCR amplified using 10 ng of WI2-3695P19 fosmid DNA, WI2-3322N15 fosmid DNA, pGL4.20-HIF1A prom plasmid DNA, WI2-1251C21 fosmid DNA, and WI2-3035P11 fosmid DNA as templates respectively. Deletion constructs were amplified from plasmids containing full length promoters. Fosmids were obtained from BACPAC genomics resource center (BACPAC Genomics, Inc. Redmond, WA) and isolated by HighPrep Plasmid DNA Kit (Magbio Genomics Inc. Gaithersburg, MD cat no. 501656596) whereas the pGL4.20-HIF1A prom plasmid was a gift from Alex Minella (Addgene plasmid # 40173 ; http://n2t.net/addgene:40173 ; RRID:Addgene_40173) and isolated by Zyppy Plasmid Miniprep Kit (Zymo Research Irvine, CA cat no. D4020).. All PCRs were performed using LongAmp Taq DNA polymerase (NEB, cat no. 50994936) in a 25 μL reaction volume according to manufacturer’s protocol. Resulting amplicons were purified by gel extraction using Wizard SV Gel and PCR Clean-Up System (Promega cat no. PR-A9281) then cloned into TOPO TA pcR2.1 (Invitrogen cat no. 45-064-1). All primers utilized to generate each construct are listed in **Supplementary Table 5**. Of note, nested PCRs were ultimately necessary to clone the GOLGA3 promoter and chr12 LG4. For constructs amplified by nested PCR, the primers listed in **Supplementary Table 5** are numbered to indicate reaction order. All resulting clones were verified by sequencing (Eurofins USA Lancaster, PA).

---

**Supplemental Figure 1. EQuIP (Enhancer Quadruplex Immuno Precipitation) PCR full gels.** PCRs employing DNA template isolated from PC3 cell EQuIP pulldowns or total input DNA collected prior to IP. PCR amplicons (∼300bp) located: (Chr5 LG4) within the Chr5 LG4, (MYO10 Prom) ∼200 bp upstream of the transcription start site of the MYO10 protein coding gene, (EXOC3 Prom1kb) ∼1.4 kbp upstream of the transcription start site of the EXOC3 protein coding gene, (EXOC3 Prom2kb) ∼2.0 kbp upstream of the transcription start site of the EXOC3 protein coding gene, (AHRR 3’ Intron) within an intron located near the 3’ end of the AHRR protein coding gene. PCR amplicons were verified by sequencing.

**Supplemental Figure 2. Relative sizes of EXOC3 and GOLGA3 promoter and Chr5 and Chr12 LG4 enhancer sequences.** Images of 1.5% agarose Tris-glycine gels ran at 4°C for 8 h at 75 V and then stained for 24 h with SYBR GOLD. Each construct was run on a gel in either the unfolded (H2O) or folded (KCl) state as indicated above the image. Scram: EXOC3-size-matched, scrambled control. s including more than one construct were folded together. The red arrow denotes the gel shift observed when the EXOC3 promoter positive strand and LG4 negative strand are folded together. 7kb ssDNA Ladder (Revvity cat# CLS157950) has 6 size standards at 1100 nt, 2100 nt, 3200 nt, 4000 nt, 5100 nt and 7200 nt.

**Supplemental Figure 3. EXOC3 promoter deletion construct sequences.** Construct names correspond to the regions indicated in Figure 5A. Regions in the 5’ half and 3’ half constructs overlapping the Middle 300bp construct are only included in the Middle 300bp construct. ≥3 consecutive genomic Gs shown in red and ≥3 consecutive genomic Cs shown in blue.

**Supplemental Figure 4. Expressions of genes regulated by LG4 enhancers are coregulated. (A)** Screen capture of UALCAN entry^SF2(1),SF2(2)^ for the top 5 genes positively correlated with EXOC3 in lung adenocarcinoma (LUAD) TCGA paient samples. **(B)** Chr5 LG4 target gene expressions in available NCBI GEO datasets taken from two distinct isolates of human lung (A549), breast (MCF7), prostate (PC3) and kidney (HEK293) cell lines as determined by standard Borchert Lab protocols ^SF2(3)^. **(C)** Screen capture of GTEx Multi Gene Query ^SF2(4)^. (**D**) Screen capture of UALCAN entry for the top 5 genes positively correlated with GOLGA3 in sarcoma (SARC) TCGA paient samples.

**Supplemental Table 1. GGG count 5kb upstream of all human mRNA promoters.**

**Supplemental Table 2. Chr5 LG4 SNPs perturbing EXOC3 expression.**

**Supplemental Table 3. Select Pore-C reads containing segments of both an LG4 and its putative target promoter.**

**Supplemental Table 4. Select Pore-C read genomic alignments.**

**Supplemental Table 5. Oligonucleotide master list.**

## References

1. Razin, S. V., Ulianov, S. V. & Iarovaia, O. V. Enhancer Function in the 3D Genome. Genes (Basel). 14, (2023).

2. Furlong, E. E. M. & Levine, M. Developmental enhancers and chromosome topology. Science 361, 1341–1345 (2018).

3. Ptashne, M. Gene regulation by proteins acting nearby and at a distance. Nature 322, 697–701 (1986).

4. Dekker, J., Rippe, K., Dekker, M. & Kleckner, N. Capturing chromosome conformation. Science 295, 1306–1311 (2002).

5. Golov, A. K., Gavrilov, A. A., Kaplan, N. & Razin, S. V. A genome-wide nucleosome-resolution map of promoter-centered interactions in human cells corroborates the enhancer-promoter looping model. doi:10.1101/2023.02.12.528105v1

6. Symmons, O. et al. Functional and topological characteristics of mammalian regulatory domains. Genome Res. 24, 390–400 (2014).

7. Kaiser, V. B. & Semple, C. A. When TADs go bad: Chromatin structure and nuclear organisation in human disease. F1000Research 6, (2017).

8. Razin, S. V. & Ulianov, S. V. Gene functioning and storage within a folded genome. Cell. Mol. Biol. Lett. 22, (2017).

9. Valton, A. L. & Dekker, J. TAD disruption as oncogenic driver. Curr. Opin. Genet. Dev. 36, 34– 40 (2016).

10. Sanyal, A., Lajoie, B. R., Jain, G. & Dekker, J. The long-range interaction landscape of gene promoters. Nature 489, 109–113 (2012).

11. Schoenfelder, S. & Fraser, P. Long-range enhancer-promoter contacts in gene expression control. Nat. Rev. Genet. 20, 437–455 (2019).

12. Deaton, A. M. & Bird, A. CpG islands and the regulation of transcription. Genes Dev. 25, 1010– 1022 (2011).

13. Huppert, J. L. & Balasubramanian, S. G-quadruplexes in promoters throughout the human genome. Nucleic Acids Res. 35, 406–413 (2007).

14. Lago, S. et al. Promoter G-quadruplexes and transcription factors cooperate to shape the cell type-specific transcriptome. Nat. Commun. 12, (2021).

15. Hegyi, H. Enhancer-promoter interaction facilitated by transiently forming G-quadruplexes. Sci. Rep. 5, 9165 (2015).

16. Burge, S., Parkinson, G. N., Hazel, P., Todd, A. K. & Neidle, S. Quadruplex DNA: sequence, topology and structure.1 S. Burge, G. N. Parkinson, P. Hazel, A. K. Todd and S. Neidle, Nucleic Acids Res., 2006, 34, 5402–15. Nucleic Acids Res. 34, 5402–15 (2006).

17. Bochman, M. L., Paeschke, K. & Zakian, V. A. DNA secondary structures: Stability and function of G-quadruplex structures. Nature Reviews Genetics 13, 770–780 (2012).

18. Huppert, J. L. & Balasubramanian, S. Prevalence of quadruplexes in the human genome. Nucleic Acids Res. 33, 2908–2916 (2005).

19. Todd, A. K., Johnston, M. & Neidle, S. Highly prevalent putative quadruplex sequence motifs in human DNA. Nucleic Acids Res. 33, 2901–2907 (2005).

20. Chambers, V. S. et al. High-throughput sequencing of DNA G-quadruplex structures in the human genome. Nat. Biotechnol. 33, 877–881 (2015).

21. Larson, E. D., Duquette, M. L., Cummings, W. J., Streiff, R. J. & Maizels, N. MutSalpha binds to and promotes synapsis of transcriptionally activated immunoglobulin switch regions. Curr. Biol. 15, 470–4 (2005).

22. Maizels, N. Dynamic roles for G4 DNA in the biology of eukaryotic cells. Nature Structural and Molecular Biology 13, 1055–1059 (2006).

23. Williams, J. D. et al. Characterization of long G4-rich enhancer-associated genomic regions engaging in a novel loop:loop ‘G4 Kissing’ interaction. Nucleic Acids Res. 48, 5907–5925 (2020).

24. Zerbino, D. R., Wilder, S. P., Johnson, N., Juettemann, T. & Flicek, P. R. The ensembl regulatory build. Genome Biol. 16, 56 (2015).

25. Cunningham, F. et al. Ensembl 2015. Nucleic Acids Res. 43, D662–9 (2015).

26. Leinonen, R., Sugawara, H. & Shumway, M. The sequence read archive. Nucleic Acids Res. 39, (2011).

27. Kodama, Y., Shumway, M., Leinonen, R. & International Nucleotide Sequence Database, C. The Sequence Read Archive: explosive growth of sequencing data. Nucleic Acids Res 40, D54–6 (2012).

28. Kubo, N. et al. Promoter-proximal CTCF binding promotes distal enhancer-dependent gene activation. Nat. Struct. Mol. Biol. 28, 152–161 (2021).

29. Fishilevich, S. et al. GeneHancer: genome-wide integration of enhancers and target genes in GeneCards. Database (Oxford). 2017, (2017).

30. Rosenbloom, K. R. et al. ENCODE data in the UCSC Genome Browser: year 5 update. Nucleic Acids Res. 41, (2013).

31. Jiang, Y. et al. SEdb: A comprehensive human super-enhancer database. Nucleic Acids Res. 47, D235–D243 (2019).

32. Zhou, Y. H. et al. Genetic Modifiers of Cystic Fibrosis Lung Disease Severity: Whole-Genome Analysis of 7,840 Patients. Am. J. Respir. Crit. Care Med. 207, 1324–1333 (2023).

33. Aguet, F. et al. Genetic effects on gene expression across human tissues. Nature 550, 204–213 (2017).

34. Deshpande, A. S. et al. Identifying synergistic high-order 3D chromatin conformations from genome-scale nanopore concatemer sequencing. Nat. Biotechnol. 40, 1488–1499 (2022).

35. Gridina, M. & Fishman, V. Multilevel view on chromatin architecture alterations in cancer. Front. Genet. 13, (2022).

36. Kim, P. et al. FusionGDB 2.0: fusion gene annotation updates aided by deep learning. Nucleic Acids Res. 50, D1221–D1230 (2022).

37. Borchert, G. M., Holton, N. W., Edwards, K. A., Vogel, L. A. & Larson, E. D. Histone H2A and H2B Are Monoubiquitinated at AID-Targeted Loci. PLoS One 5, e11641 (2010).

38. De Armond, R., Wood, S., Sun, D., Hurley, L. H. & Ebbinghaus, S. W. Evidence for the presence of a guanine quadruplex forming region within a polypurine tract of the hypoxia inducible factor 1alpha promoter. Biochemistry 44, 16341–16350 (2005).

39. Li, A. Z., Huang, H., Re, X., Qi, L. J. & Marx, K. A. A gel electrophoresis study of the competitive effects of monovalent counterion on the extent of divalent counterions binding to DNA. Biophys. J. 74, 964–973 (1998).

40. Hänsel-Hertsch, R. et al. G-quadruplex structures mark human regulatory chromatin. Nat. Genet. >48, 1267–1272 (2016).

41. Hänsel-Hertsch, R., Spiegel, J., Marsico, G., Tannahill, D. & Balasubramanian, S. Genome-wide mapping of endogenous G-quadruplex DNA structures by chromatin immunoprecipitation and high-throughput sequencing. Nat. Protoc. 13, 551–564 (2018).

42. Lyu, J., Shao, R., Kwong Yung, P. Y. & Elsässer, S. J. Genome-wide mapping of G-quadruplex structures with CUT&Tag. Nucleic Acids Res. 50, E13 (2022).

43. Hou, Y. et al. Integrative characterization of G-Quadruplexes in the three-dimensional chromatin structure. Epigenetics 14, 894–911 (2019).

44. Gibcus, J. H. & Dekker, J. The Hierarchy of the 3D Genome. Mol. Cell 49, 773–782 (2013).

45. Cremer, T. & Cremer, M. Chromosome territories. Cold Spring Harb. Perspect. Biol. 2, (2010).

46. Dixon, J. R. et al. Topological domains in mammalian genomes identified by analysis of chromatin interactions. Nature 485, 376–380 (2012).

47. Sun, F. et al. Promoter-Enhancer Communication Occurs Primarily within Insulated Neighborhoods. Mol. Cell 73, 250–263.e5 (2019).

48. Dowen, J. M. et al. Control of cell identity genes occurs in insulated neighborhoods in mammalian chromosomes. Cell 159, 374–387 (2014).

49. Hnisz, D., Day, D. S. & Young, R. A. Insulated Neighborhoods: Structural and Functional Units of Mammalian Gene Control. Cell 167, 1188–1200 (2016).

50. Hnisz, D. et al. Activation of proto-oncogenes by disruption of chromosome neighborhoods. Science (80-.). 351, 1454–1458 (2016).

51. Lupiáñez, D. G. et al. Disruptions of topological chromatin domains cause pathogenic rewiring of gene-enhancer interactions. Cell 161, 1012–1025 (2015).

52. Eddy, J. & Maizels, N. Conserved elements with potential to form polymorphic G-quadruplex structures in the first intron of human genes. Nucleic Acids Res. 36, 1321–1333 (2008).

53. Tikhonova, P. et al. DNA G-Quadruplexes Contribute to CTCF Recruitment. Int. J. Mol. Sci. 22, (2021).

54. Merkenschlager, M. & Nora, E. P. CTCF and Cohesin in Genome Folding and Transcriptional Gene Regulation. Annu. Rev. Genomics Hum. Genet. 17, 17–43 (2016).

55. Weintraub, A. S. et al. YY1 Is a Structural Regulator of Enhancer-Promoter Loops. Cell 171, 1573–1588.e28 (2017).

56. Li, L. et al. YY1 interacts with guanine quadruplexes to regulate DNA looping and gene expression. Nat. Chem. Biol. 17, 161–168 (2021).

57. Roy, S. S. et al. Artificially inserted G-quadruplex DNA secondary structures induce long-distance chromatin activation. doi:10.1101/2023.11.27.568900

58. Chandrashekar, D. S. et al. UALCAN: An update to the integrated cancer data analysis platform. Neoplasia 25, 18–27 (2022).

59. Dang, H. et al. Mining GWAS and eQTL data for CF lung disease modifiers by gene expression imputation. PLoS One 15, (2020).

60. Devadoss, D. et al. A long noncoding RNA antisense to ICAM-1 is involved in allergic asthma associated hyperreactive response of airway epithelial cells. Mucosal Immunol. 14, 630–639 (2021).

61. Devadoss, D. et al. Long noncoding transcriptome in chronic obstructive pulmonary disease. American Journal of Respiratory Cell and Molecular Biology 61, 678–688 (2019).

62. Devadoss, D. et al. Distinct Mucoinflammatory Phenotype and the Immunomodulatory Long Noncoding Transcripts Associated with SARS-CoV-2 Airway Infection. medRxiv Prepr. Serv. Heal. Sci. (2021). doi:10.1101/2021.05.13.21257152

63. Devadoss, D., et al. Immunomodulatory LncRNA on antisense strand of ICAM-1 augments SARS-CoV-2 infection-associated airway mucoinflammatory phenotype. iScience 25, (2022).

64. Manevski, M. et al. Increased Expression of LASI lncRNA Regulates the Cigarette Smoke and COPD Associated Airway Inflammation and Mucous Cell Hyperplasia. Front. Immunol. 13, (2022).

65. Wang, Y. et al. SEdb 2.0: a comprehensive super-enhancer database of human and mouse. Nucleic Acids Res. 51, D280–D290 (2023).

66. Altschul, S. F., Gish, W., Miller, W., Myers, E. W. & Lipman, D. J. Basic local alignment search tool. J. Mol. Biol. 215, 403–410 (1990).

