## Supplementary figures and images for "Long G4-rich enhancer physically interacts with EXOC3 promoter via a G4:G4 DNA-based mechanism"

### Supplemental Figure 1

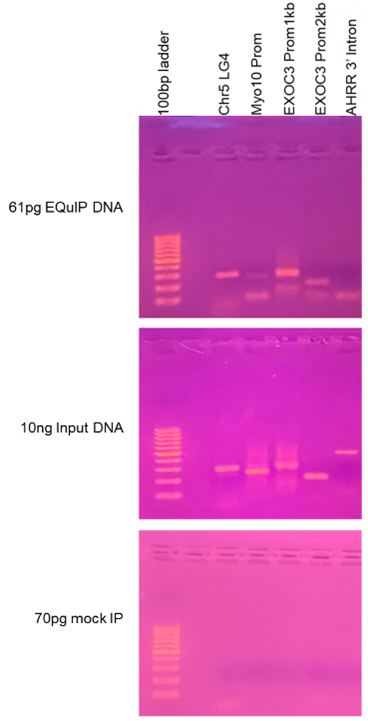

### Supplemental Figure 2

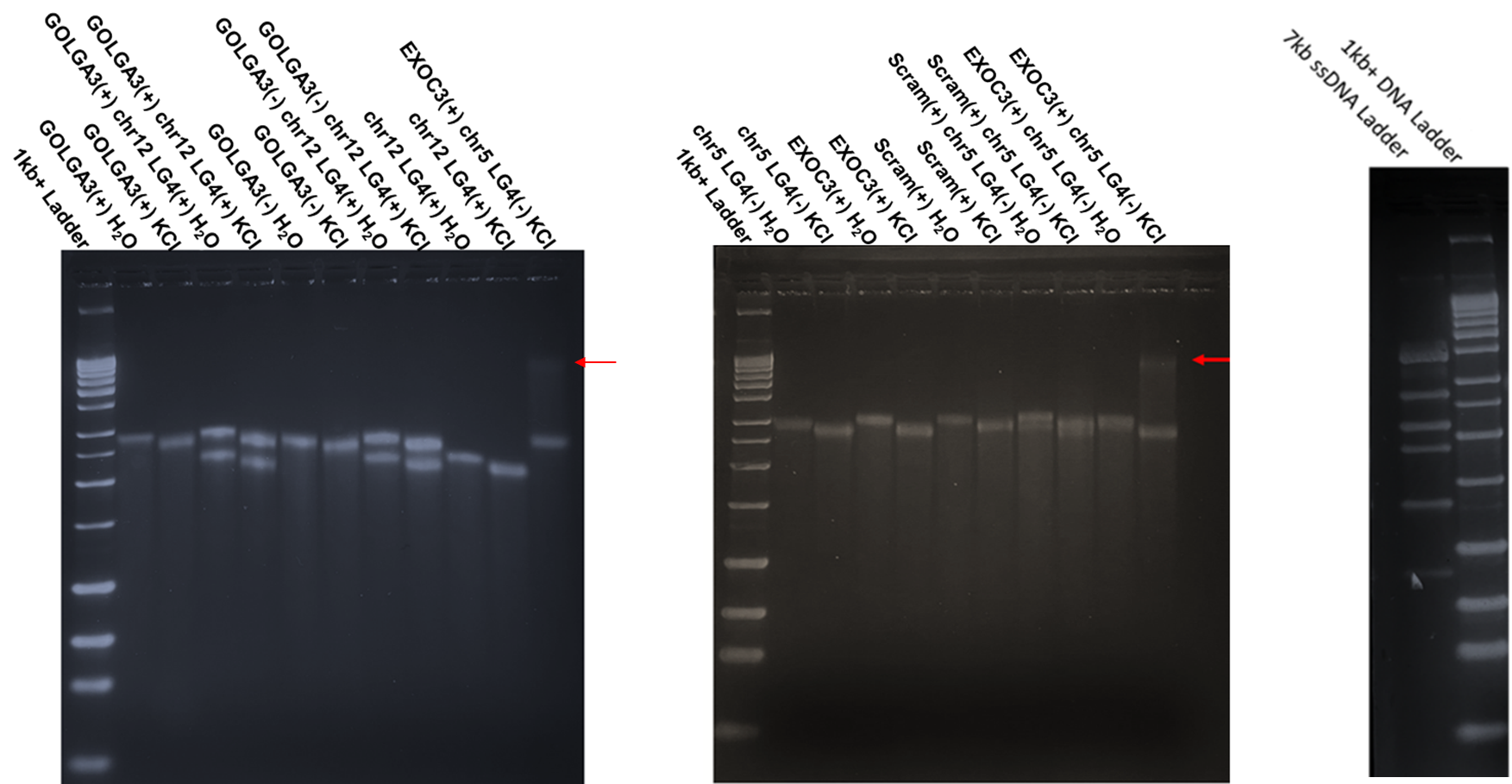

### Supplemental Figure 3

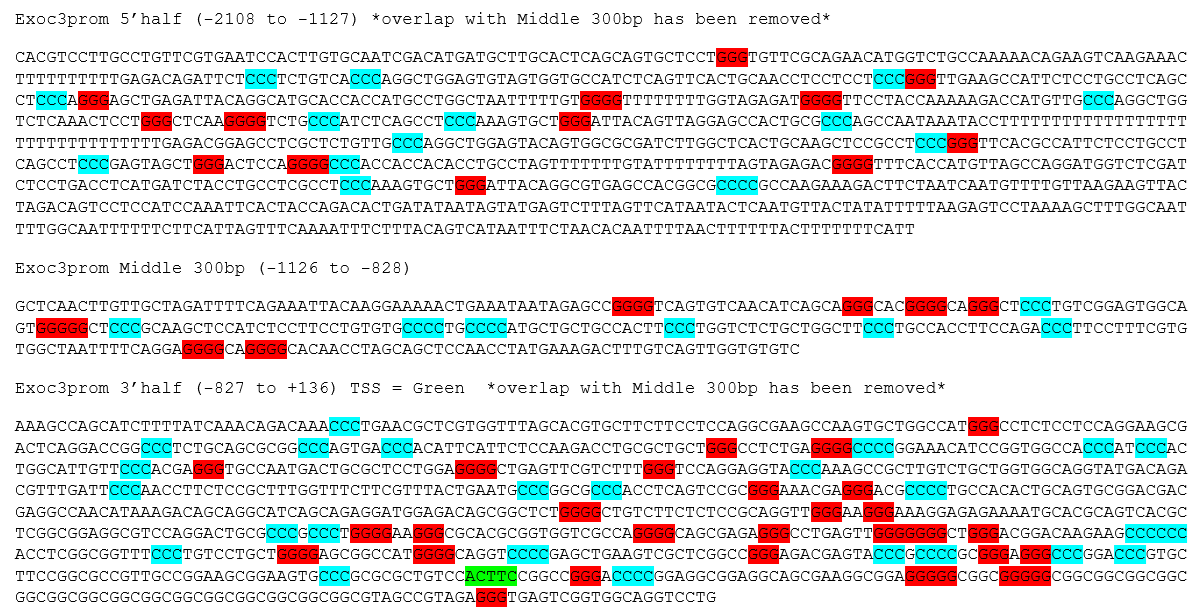

### Supplemental Figure 4

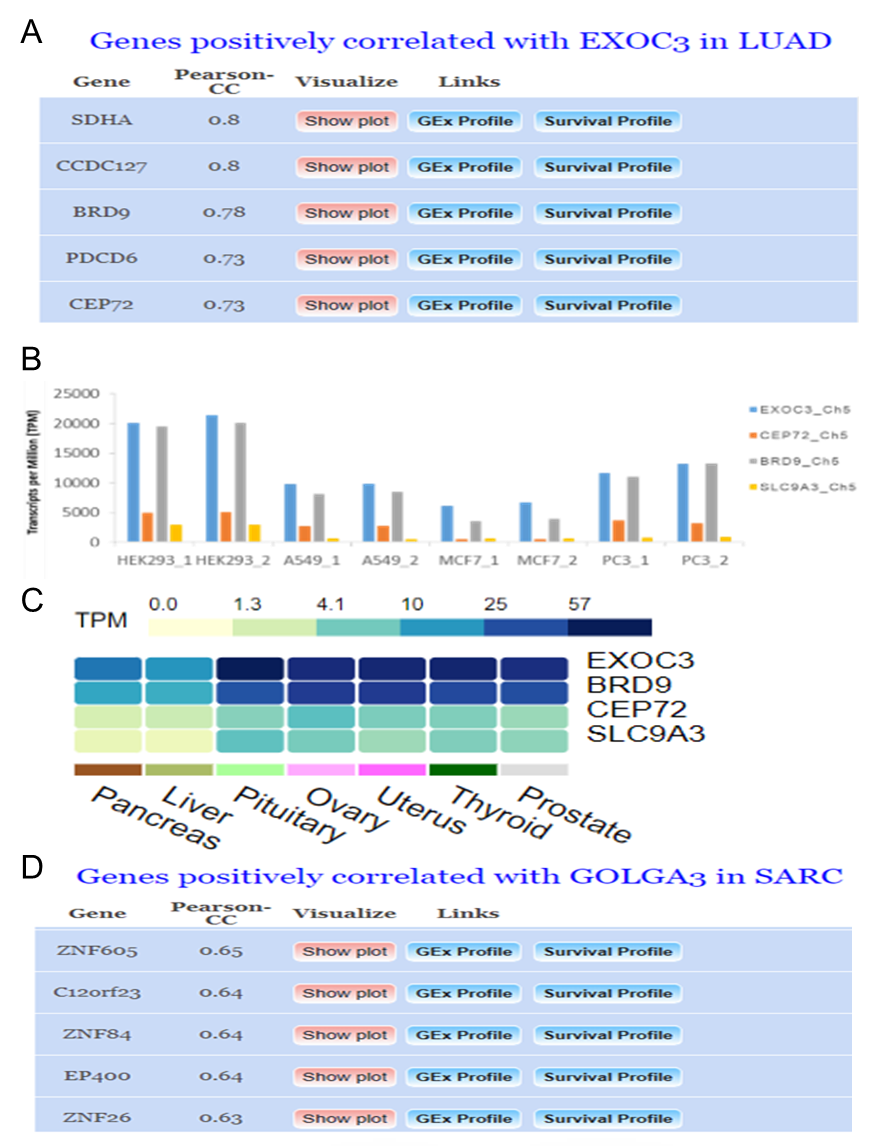

### Supplemental Table 1

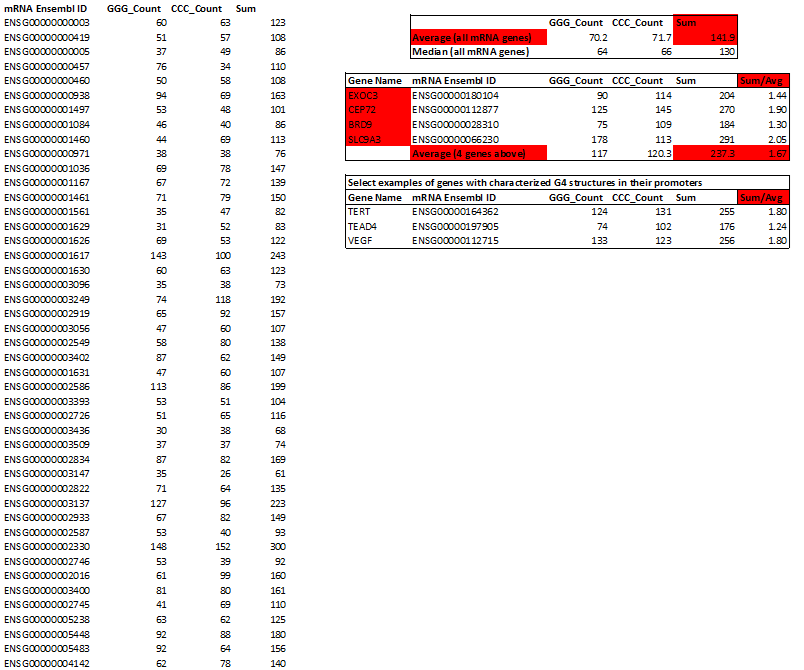

### Supplemental Table 2

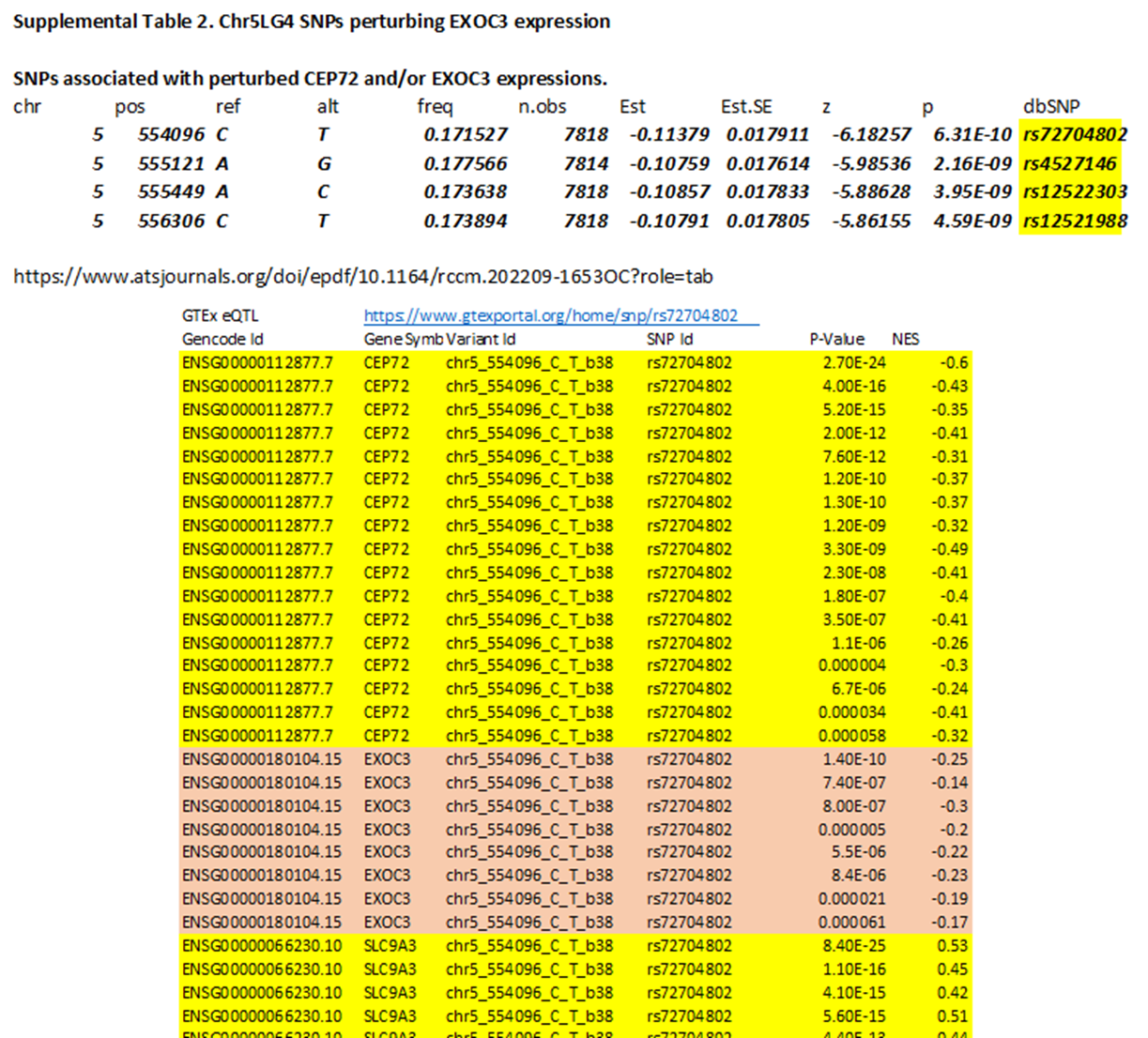

### Supplemental Table 3

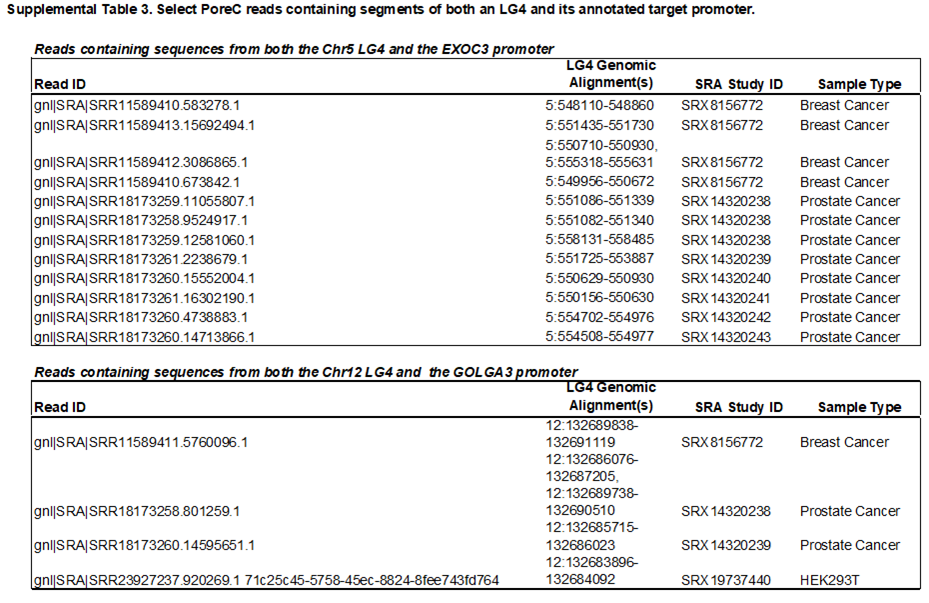

### Supplemental Table 4

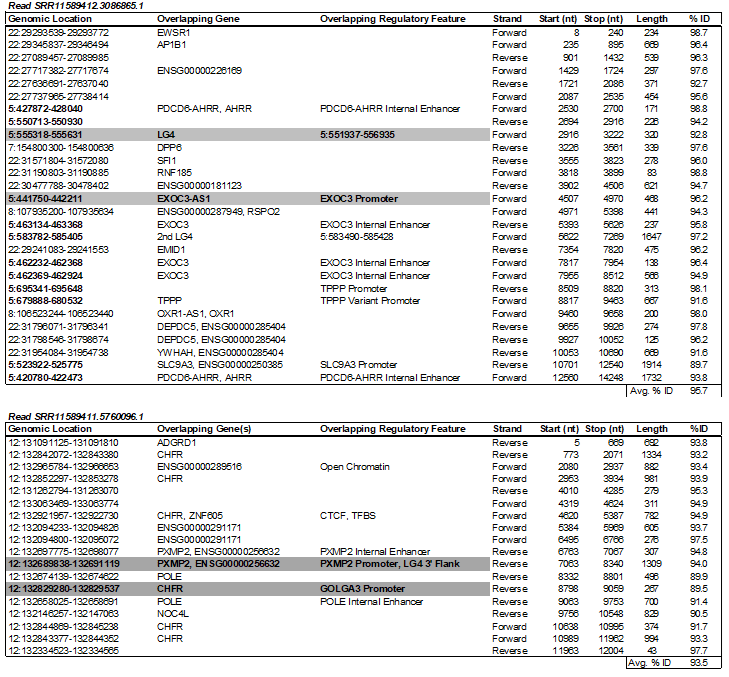

### Supplemental Table 5

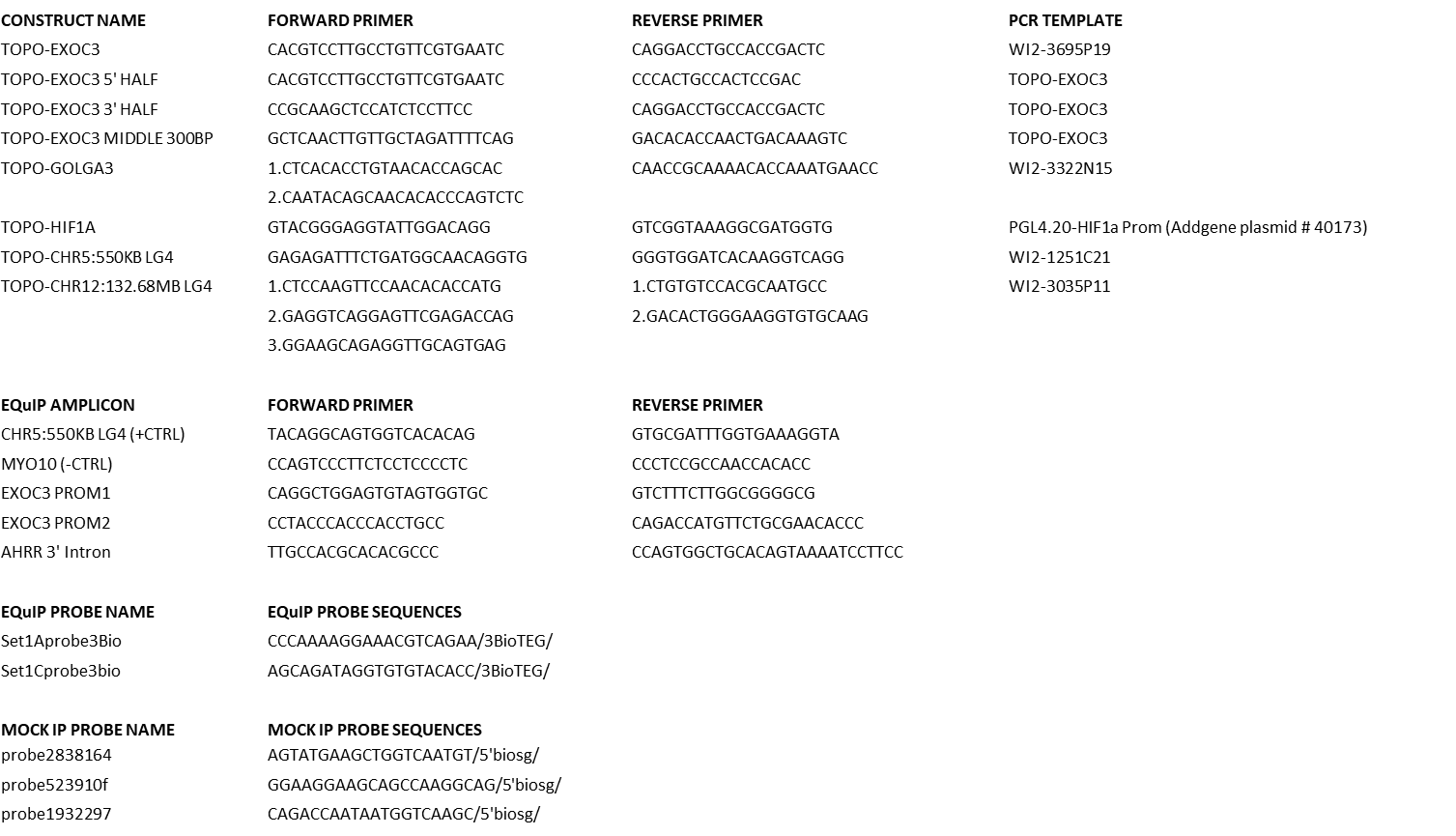
